# Traffic Jam activates the *Flamenco* piRNA cluster locus and the Piwi pathway to ensure transposon silencing and *Drosophila* fertility

**DOI:** 10.1101/2024.08.15.608167

**Authors:** Austin Rivera, Jou-Hsuan Roxie Lee, Shruti Gupta, Linda Yang, Raghuveera Kumar Goel, Joseph Zaia, Nelson C. Lau

## Abstract

*Flamenco (Flam)* is the most prominent piRNA cluster locus expressed in *Drosophila* ovarian follicle cells, and it is required for female fertility to silence *gypsy/mdg4* transposons. To determine how *Flam* is regulated, we used promoter-bashing reporter assays in OSS cells to uncover novel enhancer sequences within the first exons of *Flam*. We confirmed the enhancer sequence relevance in vivo with new *Drosophila Flam* deletion mutants of these regions that compromised *Flam* piRNA expression and lowered female fertility from activated transposons. Our proteomic analysis of proteins associated with these enhancer sequences discovered the transcription factor Traffic Jam (TJ). *Tj* knockdowns in OSS cells caused a decrease in *Flam* transcripts, *Flam* piRNAs, and multiple Piwi pathway genes. A TJ ChIP-seq analysis from whole flies and OSS cells confirmed TJ binding exactly at the enhancer that was deleted in the new *Flam* mutant as well as at multiple Piwi pathway gene enhancers. Interestingly, TJ also bound the Long Terminal Repeats of transposons that had decreased expression after *Tj* knockdowns in OSS cells. Our study reveals the integral role TJ plays in the on-going arms race between selfish transposons and their suppression by the host Piwi pathway and the *Flam* piRNA cluster locus.

## INTRODUCTION

In the *Drosophila* ovary, the germline compartment of nurse cells and the oocyte are encompassed by the somatic compartment of follicle cells. The germline generates Piwi-interacting RNAs (piRNAs) from one set of piRNA cluster loci that express transcripts from both genomic strands and are transcriptionally activated by the factors Moonshiner, TRF2 and the Rhino-Deadlock-Cutoff complex (Mohn et al. 2014; Andersen et al. 2017). In contrast, follicle cells only express a distinct set of uni-strand piRNA cluster loci, with the most prominent one named *Flamenco* (*Flam*) for the initial genetic discovery of mutants in this locus that enabled the mobilization of the *gypsy/mdg4* Transposable Elements (TEs) (Robert et al. 2001; Sarot et al. 2004; Brennecke et al. 2007).

Although *Drosophila* mutants deleted of individual germline dual-strand piRNA cluster loci are nearly as fertile as wildtype flies with little TE activation (Gebert et al. 2021), *Flam* mutant females are severely subfertile because multiple retroviral TEs are activated and generate infectious viral particles that are linked to massive ovarian collapse (Pelisson et al. 1994; Prud’homme et al. 1995; Mevel-Ninio et al. 2007). Two classic *Flam* mutations, KG00476 and BG02658, are referred to as *Flam^KG^* and *Flam^BG^* in this study and are both caused by *P-element* insertions in the *Flam* promoter/enhancer regions that result in severe loss of *Flam* piRNAs (Brennecke et al. 2007; Mevel-Ninio et al. 2007; Malone et al. 2009). A previous study proposed the *Flam^BG^* mutation disrupted a binding site for the transcription factor *Cubitus interruptus* (*Ci*) (Goriaux et al. 2014). Because *Ci* is also broadly expressed in many other *Drosophila* cell types that lack *Flam* and piRNA expression (Wang and Holmgren 1999), we wondered if another factor specific to follicle cells could be the primary activator of *Flam*.

One set of *Drosophila* cell cultures, the Ovarian Somatic Sheet (OSS) and the Ovarian Somatic Cell (OSC) lines are derived from the follicle cell lineage and express the complete PIWI pathway including abundant genic piRNAs and *Flam* piRNAs (Niki et al. 2006; Lau et al. 2009; Saito et al. 2009). These OSC/OSS lines have been important models for illuminating the somatic compartment of PIWI/piRNA biochemical silencing mechanisms and piRNA biogenesis factor roles (Haase et al. 2010; Saito et al. 2010; Sienski et al. 2012; Post et al. 2014), but germline piRNA cluster loci and germline *Aub* and *Ago3* homologs of *piwi* are not expressed in these cells. Only the loss of the *l(3)mbt* and *lint-O* repressors in OSCs transformed these cells into expressing *Aub*, *Ago3* and *vasa*, and yet the piRNAs in these modified OSCs still mainly resembled the original bulk of *Flam* piRNAs (Sumiyoshi et al. 2016; Yamamoto-Matsuda et al. 2022).

Although the high-throughput STARR-seq method could define new OSC-specific enhancer elements within the *Flam* locus (Arnold et al. 2014), neither the STARR-seq approach nor bioinformatics predictions (Goriaux et al. 2014; Rivera et al. 2019) have determined which specific enhancer elements and transcription factor(s) fully drive *Flam* expression. Therefore, we approached this knowledge gap by re-designing a promoter-bashing reporter assay for *Flam* to define enhancer sequences, and then utilized these enhancer sequences to discover the specific DNA-binding protein with proteomics. Our study discovers the large MAF transcription factor Traffic Jam (TJ) binding to *Flam* enhancers and directly regulating *Flam* expression. Additionally, we show *Tj*’s broad influence on the follicle cell transcriptome by also serving as a nexus within the arms-race conflict between the *Drosophila* host genome defense pathways and the selfish TEs.

## RESULTS

### A revised promoter-bashing reporter assay pinpoints new Flam enhancer elements

A previously described *Flam* promoter luciferase reporter assay (Goriaux et al. 2014) had first started with a construct that was 515 bp upstream and 101 bp downstream of the *Flam* transcription start site (TSS). This 616 bp *Flam* promoter element reporter tested deleting the TSS and a putative binding sequences for the *Ci* transcription factor. We also tested a very short fragment from *Flam* we designated as a negative control (segment i14) by first transfecting solely *Drosophila* OSS cells like (Goriaux et al. 2014). At first, this original segment i4 appeared to promote gene activation compared to our negative control (**Figure 1A**).

**Figure 1.**
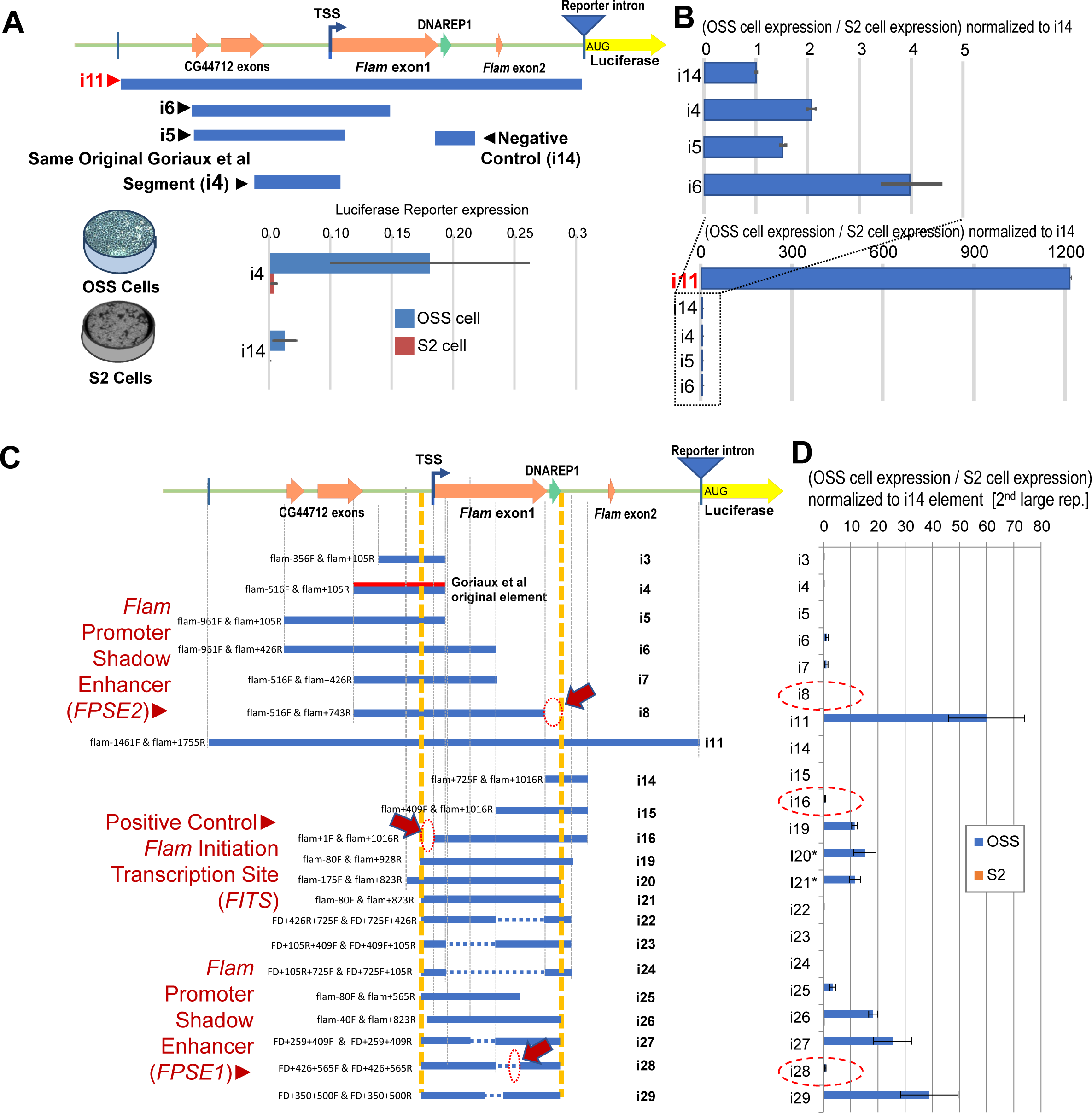
Promoter-bashing reporter assay discovers novel *Flam* enhancer elements. (A) Diagram of *Flam* promoter region targeted in a novel format of a promoter bashing assay that utilizes an intron upstream of the luciferase. (B) Snapshot of normalized reporter assay results comparing the Goriaux et al i4 element to the i11 whole *Flam* promoter region (C) Complete diagram of bashing constructs highlighting discovery of *FITS*, *FPSE1* and *FPSE2* regions. (D) Corresponding normalized promoter bashing assay results from the constructs depicted in (C). Small text labels are the names of the primers used to generate the constructs. Deletions of interest marked by dashed ovals. *The constructs i20* and i21* were first indication that expression in OSS cells required the extra reporter intron. Constructs i3–i19 had similar expression result as constructs that lacked the reporter intron, and i22–i29 were only constructed with the reporter intron.

However, to discern away from general transcription activation and to focus on OSS cell-specific activation of the *Flam* promoter reporters, we included additional reporter transfections into S2 cells that completely lacked expression of the Piwi pathway and *Flam* (Saito et al. 2006; Saito et al. 2009). We then analyzed reporter expression levels through first calculating the ratio of OSS cell reporter readings over S2 cell readings, and then normalizing these ratios of readings to the segment i14 negative control element. This analysis showed only a modest (≤2 fold) basal transcription activation difference between the Goriaux et al element i4 and other similarly sized putative *Flam* enhancer constructs (Fig. 1B). With this revised promoter bashing reporter assay format, we then determined an ‘i11’ element (∼1.5kb upstream and ∼1.8 downstream of the *Flam* TSS) that displayed a >1,200-fold activation in OSS cells versus S2 cells.

Since this i11 sequence included original intronic elements proximal to the *Flam* TSS, we controlled for RNA splicing impact by adding a minimal synthetic intron just upstream of luciferase, and this completely mitigated splicing artifacts on reporters lacking this synthetic intron (**Supplemental Figure S1A**). This revised promoter bashing reporter assay reconfirmed the importance of the *Flam* TSS in the *Flam* promoter luciferase reporter but could not recapitulate an impact for the *Ci* binding site nor a role for *Ci* in *Flam* activation (Supp. Fig. S1B). Our siRNA knockdowns of *Ci* in OSS cells also did not affect *Flam* RNA levels nor raise TE expression (Supp. Fig. S1C).

From these data, our goal was to deduce which discrete sequence elements activate the *Flam* promoter reporters most specially in OSS cells (Fig. 1C), to go beyond general transcription activation signal like a *Flam* TSS. The 80 bp around the *Flam* TSS that we named the *Flam* Initiation Transcription Site (*FITS*) did behave as positive control for the *Flam* reporter, but we did not see more direct OSS cell-specific activation by other TSS adjacent sequences like i3-to-i8 elements upstream of the TSS, or i22, i25 and i28 elements with sequences immediately downstream of the *Flam* TSS. We deduced instead two other impactful elements 426 bp and 743 bp downstream of the *Flam* TSS. We named these two ∼139 bp and ∼80 bp sequences *Flam* Promoter Shadow Enhancer #1 and #2, respectively (*FPSE1*, *FPSE2*). The *FPSE1* element is a uniquely mapping sequence in the *Drosophila* genome while *FPSE2* encompasses part of a DNAREP1 repeat and a putative start of an intron.

### New deletion mutants of Flam enhancer elements exhibit compromised fertility

To test the in vivo relevance of *FITS*, *FPSE1* and *FPSE2* elements, we searched for CRISPR/Cas9 small guide RNA (sgRNA) targeting sequences that closely flanked these loci for generating deletion mutants in *Drosophila melanogaster*. We screened and obtained some precise deletion mutants arising from DNA repair directed by single-stranded oligonucleotides (ssODNs) after sgRNAs were injected into the Cas9-expressing fly line (**Figure 2A**). Possibly due to the DNAREP1 repeat in the *FPSE2* sequence, the two *FPSE2a* and *FPSE2b* mutations were less precise because additional sequences from the ssODN were then duplicated into the sgRNA-guided deletion (**Supplemental Figure S2**). All mutants were confirmed by sequencing and subjected to two-to-four rounds of backcrossing to *w1118* before two final crosses to balance over the *FM7a* balancer chromosome.

**Figure 2.**
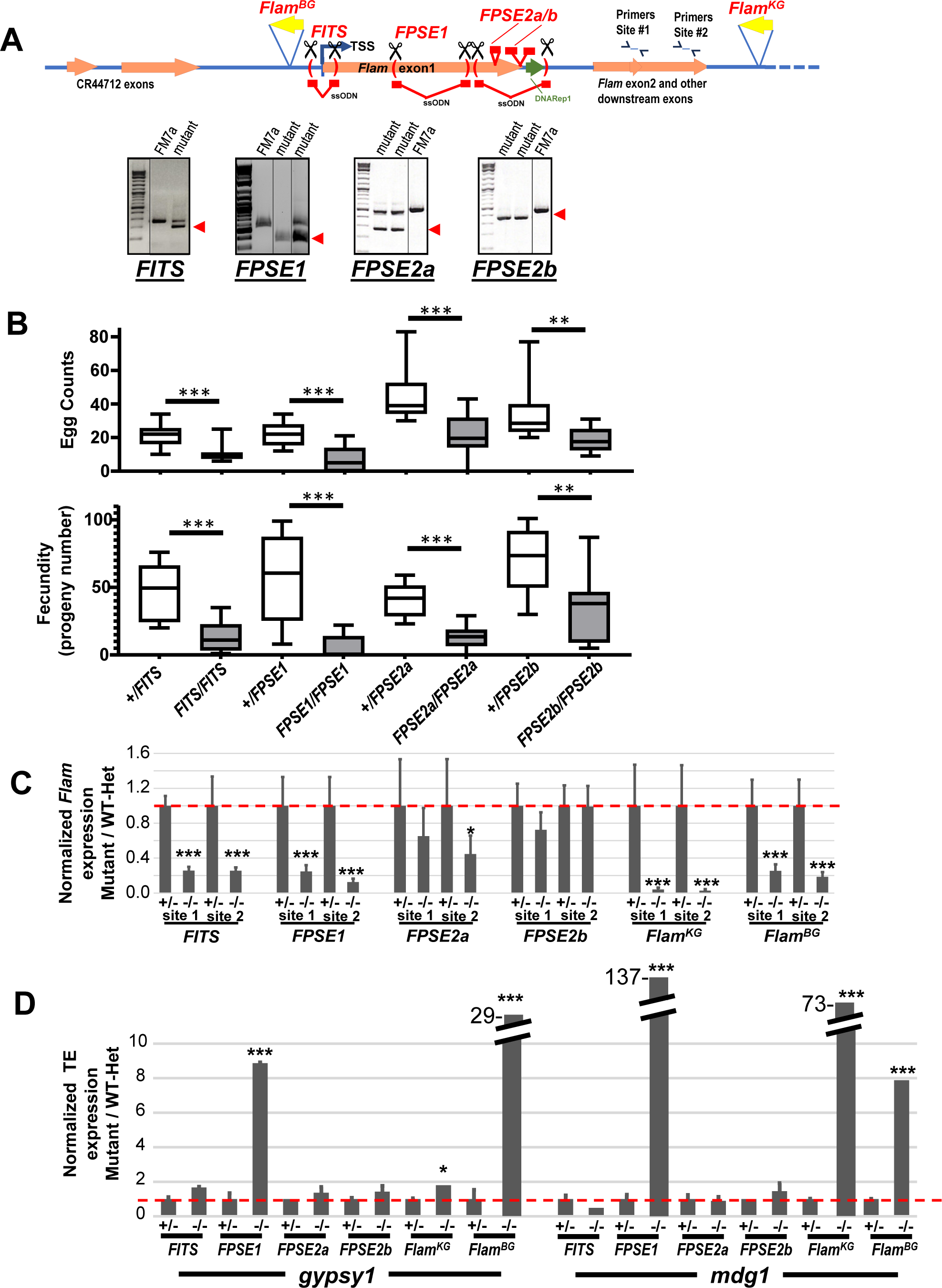
New *Flam* mutants with deletions of *FITS*, *FPSE1* and *FPSE2a*/*2b* regions have compromised female fertility. Diagram of sgRNAs and ssODN targeting of *Flam* promoter/enhancer regions. Genomic PCR showing proper locus deletions in fly mutants compared to the *FM7a* balancer line. (B) Lower female fertility and fecundity in new *Flam* promoter/enhancer region mutants. *p<0.05, **p<0.01, ***p<0.001, T-test. (C) RT-qPCR of mutants versus heterozygous siblings show *Flam* precursor RNA reduction, (D) and *gypsy1 and mdg1* RNA activation. *p<0.05, **p<0.01, ***p<0.001, Wilcoxon test.

Adult female homozygous mutants were compared to their heterozygous siblings in egg counts assays and fecundity assays (Fig. 2B). Although not completely sterile, all of the *FITS*, *FPSE1*, and *FPSE2a/b* homozygous mutants were subfertile because of the drastic reduction of mature eggs that lead to much lower fecundity. We then extracted ovarian total RNA to measure by RT-qPCR *Flam* RNA transcripts and two TE transcripts, *gypsy1* and *mdg1* known to be elevated in *Flam^KG^* mutants (Malone et al. 2009). *Flam* RNA was strongly reduced in homozygous *FITS* and FPSE1 mutants at the same severity as *Flam^BG^* and *Flam^KG^* mutants, but *Flam* RNA was much less affected in *FPSE2a/b* mutants (Fig. 2C). The strong reactivation of both *gypsy1* and *mdg1* was only observed in *Flam^BG^* and *FPSE1* homozygous mutants (Fig. 2D). Although the *FPSE1* region is downstream of the *P-element* insertion of *Flam^BG^* (Fig. 2A), these data suggest the greatest phenotypic similarity between these two mutants. These mutants’ subfertile phenotypes validated the promoter-bashing reporter assay results and also encouraged us to further characterize the long RNA and piRNA expression from the ovaries of these new *Flam* mutants.

### The FPSE1 mutation phenocopies the dramatic loss of Flam transcripts and piRNAs seen in Flam^BG^ and Flam^KG^ mutants

To expand upon the limited resolution of our RT-qPCR results, we then deeply sequenced matched long total RNA and small RNA libraries from the ovaries of homozygous *Flam* mutants, comparing relative Reads Per Million (RPM) to ovaries from heterozygous siblings. Surprisingly, despite the deletion of the wild-type *Flam* TSS, the *FITS* ovaries displayed a shift of the new TSS to a site ∼350 bp upstream of the original *Flam* TSS (**Figure 3A**). The *FITS* deletion impacted the RT-qPCR assay with primers directed at *Flam*’s exon 2, but much of the rest of the *Flam* long RNA transcript many kilobases downstream was still being efficiently expressed in *FITS* homozygous mutants (Fig. 3C). As such, *Flam* piRNAs and TE silencing and TE small RNAs were also mainly unaffected by the *FITS* mutation (Fig. 3B,C,D).

**Figure 3.**
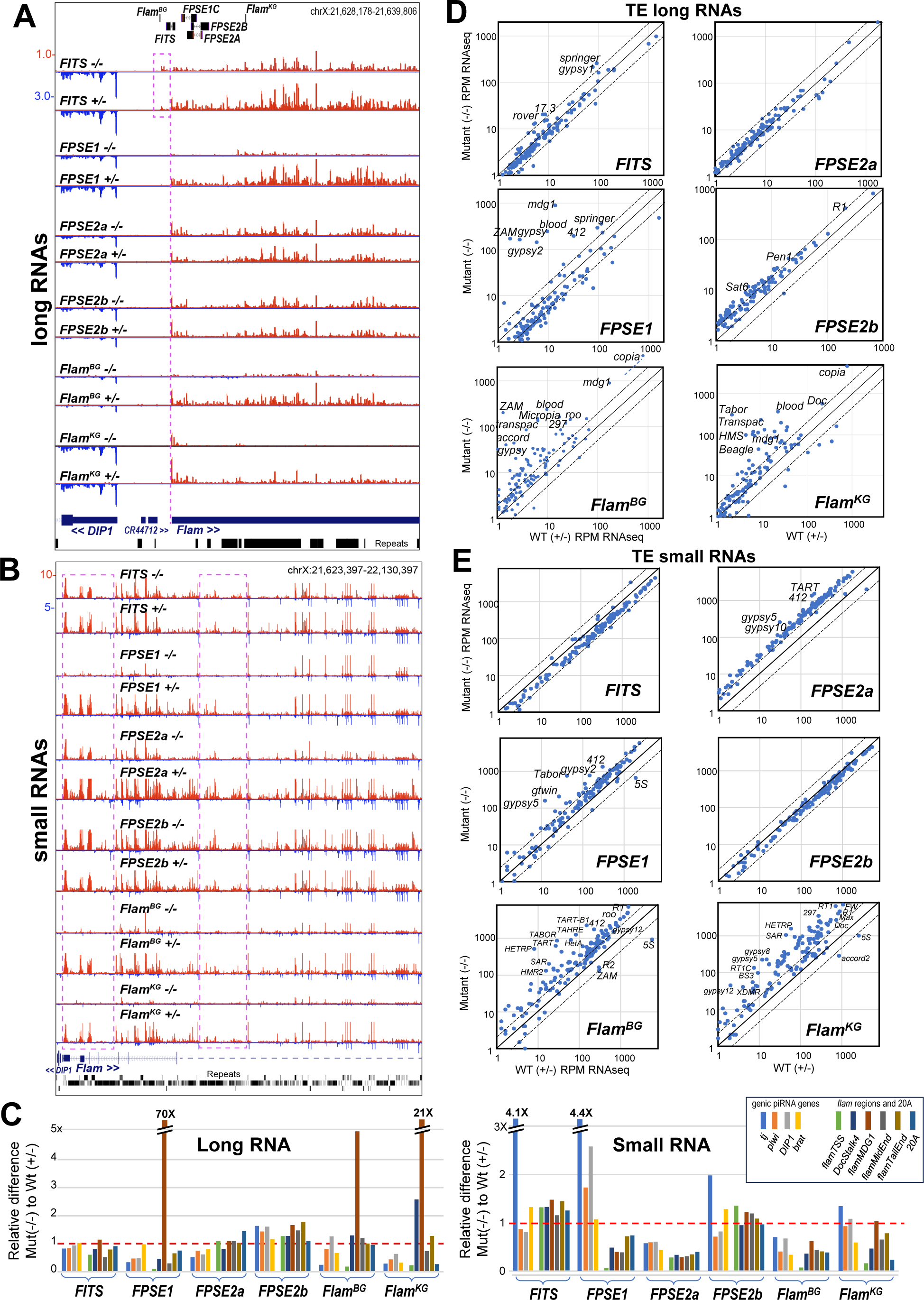
Long and small RNAseq of ovaries from the *Flam* promoter/enhancer mutants of *FITS*, *FPSE1* and *FPSE2a/2b*. (A) Genome browser snapshot of the ovary long RNAs proximal to the *Flam* Transcription Start Site (TSS) in homozygous and heterozygous mutants. The magenta dash line marks the main TSS of *Flam* precursor transcript while the magenta box highlights a new upstream TSS in *FITS* mutants that deleted to original TSS region. (B) Genome browser snapshot of the ovary small RNAs throughout the ∼500kb region of *Flam* from homozygous and heterozygous mutants. Magenta boxes mark the *flamTSS* and *flamMidEnd* regions as areas of discernable small RNA accumulation differences. (C) Quantitation of the relative difference between long RNAs and small RNAs for genic piRNA genes and *Flam* regions as well as the upstream *20A* piRNA cluster locus. Scatterplots of reads per million for TE consensus elements compared between homozygous and heterozygous mutants for (D) long RNAs and (E) small RNAs.

Although subfertile phenotypes in all the new mutant *Flam* females were reproducible (Fig. 2B), there was also no significant differences in long *Flam* transcript levels nor TE expression differences between homozygous *FPSE2a* and *FPSE2b* mutants and their heterozygous siblings (Fig. 2C,D and Fig. 3). We do not have an explanation for why phenotypes persist despite insufficient effects on *Flam* piRNAs in the *FPSE2* and *FITS* mutants, but these results indicate plasticity in the regulation of the *Flam* piRNA cluster locus and suggest the transcriptional activator binds elsewhere from the *FITS* and *FPSE2* regions.

The massive TE reactivation from loss of *Flam* long transcripts and piRNA expression first seen in *Flam^BG^* and *Flam^KG^* was recapitulated in the *FPSE1* homozygous mutants, such as the activation of *ZAM*, *blood* and *gypsy* TEs (Fig. 3D). Although *Flam* piRNAs are greatly diminished in *FPSE1* mutants as well as in *Flam^BG^* and *Flam^KB^* mutants, the small RNA libraries from ovaries also showed elevated TE piRNAs that could have come from the germline compartment that now dominated the compromised somatic compartment in these mutants (Fig. 3E).

### Proteomics discovery of Traffic Jam as a prime candidate binding Flam enhancer elements

A goal of the promoter-bashing reporter assay was to define small DNA sequences of *Flam* enhancers to then perform an unbiased biochemical discovery of the transcription factor binding to these elements. We generated 280 bp, 647 bp and 501 bp biotin-tagged PCR-amplified dsDNAs representing *FITS*, *FPSE1* and *FPSE2*, respectively, and then incubated these DNAs in OSS cell and S2 cell nuclear extracts before pulling the DNAs down with streptavidin-coupled Dynabeads. Pulled-down DNA binding proteins were then visualized with silver-stained gels and subjected to on-bead trypsin digestion and TMT-labeling for quantitative proteomic analysis (**Supplemental Figure S3A**).

Three analyses of the proteomic profiles between the different DNA elements congruently pointed to the Traffic Jam (TJ) transcription factor as the top enriched protein in *FPSE1* and *FPSE2* pulldowns (Supp. Fig. S3B,C). The *Tj* gene is known to be a critical gene in *Drosophila* follicle cell development and ovary function, and *Tj* null mutants are completely sterile from totally collapsed ovaries and testes (Li et al. 2003; Gunawan et al. 2013; Panchal et al. 2017). The *Tj* mRNA serves as a precursor for abundant genic 3’ UTR piRNAs (Robine et al. 2009; Saito et al. 2009), while also implicated to bind to the *piwi* gene promoter (Saito et al. 2009). However, the full extent of TJ’s regulation of the follicle cell transcriptome was still unknown.

We first tried to validate TJ’s potential impact on *Flam* expression in vivo with follicle-cell specific RNAi knockdown in the fly ovary starting with the *Tj-GAL-4* driver coupled with a *GAL-80^TS^* repressor for temperature-controlled triggering of the *Tj* knockdown. When *Tj* knockdown was triggered efficiently, ovaries were collapsed to rudimentary stubs lacking tissue for analysis, whereas timed *Tj* knockdowns were insufficient when ovaries were allowed to develop enough follicle cells for analysis (**Supplemental Figure S4A**). Additional later-stage follicle cell drivers like *CG13083-GAL4* also did not cause sufficient decrease of *Tj* despite allowing for ovary development (Supp. Fig. S4B). Given the very abundant expression and critical importance of *Tj* in follicle cell and ovary development, in vivo RNAi knockdown of *Tj* to study *Flam* expression remains a challenge.

To add to our validation efforts, we raised a new rabbit polyclonal antibody to the N-terminal peptide of TJ, and we determined the antibody’s specificity to be equivalent to an original guinea pig-derived anti-TJ antibody (Li et al. 2003) (Supp. Fig. S4C). We also designed specific and efficient siRNAs to knock down *Tj* in OSS cells which can survive several days and reveal TE activation upon *piwi* knockdown (Saito et al. 2009; Haase et al. 2010; Saito et al. 2010; Post et al. 2014; Sytnikova et al. 2014; Genzor et al. 2021). Our rabbit anti-TJ antibody recognized and immunoprecipitated TJ specifically and confirmed very effective TJ protein loss during siTj siRNA knockdown in OSS cells (Suppl Fig. S4C,D).

### Tj knockdown in OSS cells validates direct regulation of Flam expression

Taking advantage of the OSS cell system for siTj knockdowns, we conducted multiple validation approaches of *Tj*’s potential regulation of *Flam*. We first retested two minimal *Flam* promoter luciferase reporter constructs that either contained or lacked the *FPSE1* sequence in OSS cells after siTj knockdown (**Figure 4A**). Compared to siGFP and siPiwi negative controls, only the i29 promoter reporter containing the *FPSE1* sequence was less activated upon siTj knockdown (Fig. 4B). We then applied the same *Flam* and TEs RT-qPCR assays we had applied to the *FPSE1* mutant fly ovaries (Fig. 2C,D) to the OSS cells after siTj knockdown. Indeed, *Flam* transcripts were reduced after siTj knockdown followed by the increase of just the *gypsy1* TE but not two other TEs that are targeted for silencing by PIWI and piRNAs in OSS cells (Fig. 4C).

**Figure 4.**
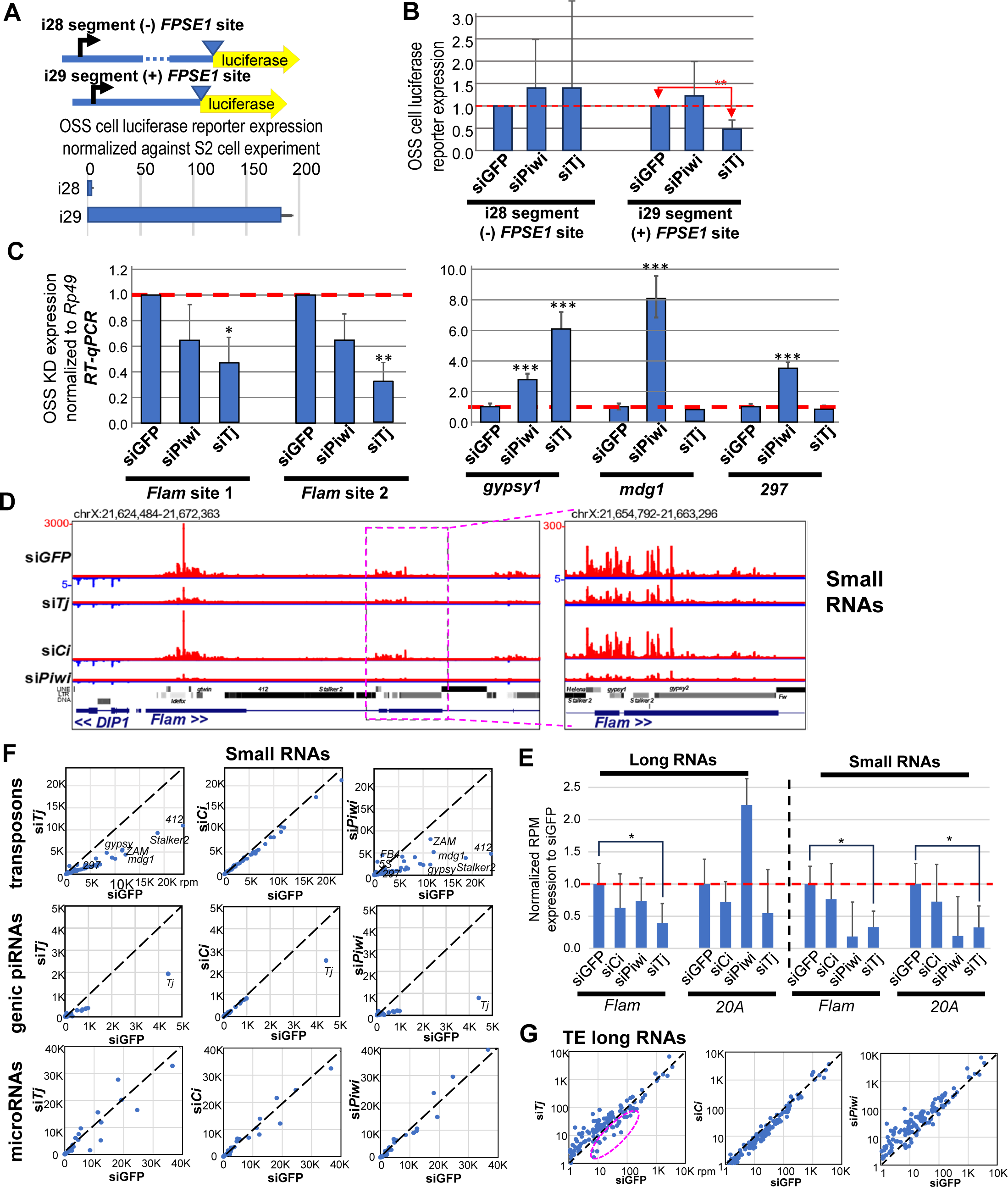
*Tj* knockdown in OSS cells decreases *Flam* promoter reporter expression, *Flam* transcripts, *Flam* piRNAs, and other piRNAs. (A) Diagram of *Flam* promoter reporters that contain or lack a putative *TJ* binding region *FPSE1* which is reflected by the respective luciferase expression levels in OSS cells relative to S2 cells. (B) Reporter assay results showing that only the ‘i29’ promoter reporter is affected negatively by the knockdown of *Tj*. (C) RT-qPCR of *Flam* precursor RNAs and transposons in OSS cells. (D) Genome browser tracks showing great reduction of *Flam* piRNAs during siTj and siPiwi knockdowns (average of three biological replicates). All tracks have the same Y-axis scale as siGFP, with a zoom inset magnified to the right. (E) Quantification of the total long and small RNAs from the *Flam* and *20A* piRNA clusters during knockdowns and through three biological replicates of deep sequencing. *<p=0.05, T-test. (F) Scatterplots comparing the changes in piRNA levels for transposon and genic piRNAs. MicroRNA changes are also shown for comparison. Linear scales chosen to better visualize differences of the averaged counts from three biological replicates. (G) Scatterplots of the averaged counts of long RNAs mapped to TEs. The dashed magenta oval notes a set of TEs with decreased long RNAs immediately after siTj knockdown.

We then sequenced long RNAs and small RNAs from OSS cells transfected 48 hours prior with siTJ, siCi, and siPiwi siRNAs to knock down target genes and compared to the siGFP control. The only piRNA effect of the siCi knockdown was a decrease in genic *Tj* piRNAs (Fig. 4F). The *Flam* piRNAs were as severely depleted in siTj compared to siPiwi, while the *Flam* long RNA was only reduced by siTj (Fig. 4D,E and **Supplemental Figure S5B**). The siTJ knockdown also reduced piRNAs from the *20A* piRNA cluster locus without a significant decrease of *20A* long RNAs (Fig. 4E). Many other TE-targeting and genic piRNAs were depleted as severely in the siTJ knockdown as the siPiwi knockdown (Fig. 4F). But in contrast to the expected observation of many elevated TE long RNAs observed in the siPiwi knockdown, a notable number of TE long RNAs had decreased in expression in the siTj knockdown (Fig. 4G).

Turning to the protein coding genes, many more genes were upregulated after siTj and siPiwi knockdown compared to the siGFP control (Supp. Fig. S5C). We inspected more closely genes downregulated after siTj as potential direct targets of transcriptional activation by *Tj*, and we observed a ∼75% decrease in *piwi* mRNA levels after siTj knockdown (Supp. Fig. S5D), which agrees with the previously described binding of TJ to the *piwi* promoter (Saito et al. 2009). Interestingly, several other piRNA biogenesis factor genes were also strongly downregulated by siTj knockdown, such as *armi*, *mael*, *fs(1)Yb* (Supp. Fig. S5D). We confirmed the proteins from these genes including PIWI were decreased in OSS cells during two independent siRNAs knocking down *Tj* (Supp. Fig. S4D). Since siTj knockdown also reduces expression of multiple Piwi pathway genes, this explains why all the OSS cell piRNAs are as severely depleted in siTj as siPiwi knockdowns (Fig. 4F).

### TJ ChIPseq reveals direct binding to Flam FPSE1 region and Piwi pathway gene promoters

To determine how directly TJ binds to target gene promoters to influence transcription, we conducted TJ ChIPseq in OSS cells as well as profiled the gene-activation H3K4me3 and gene-silencing H3K9me3 chromatin marks by Cut&Run in normal OSS cells and after siTj knockdown. We also integrated into our analysis a modENCODE ChIPseq dataset of transgenically expressed GFP-Tj from whole flies (Kudron et al. 2018), which has the caveat of the anti-GFP antibody accessing the GFP-TJ fusion from tissues other than the ovary. Nevertheless, we could find TJ ChIPseq peaks confidently from both datasets with the Irreproducibility Discovery Rate (IDR) approach and then we annotated the peaks to genes, finding 1275 genes with TJ peaks overlapping both datasets (**Figure 5A, Table S1 and S2**).

**Figure 5.**
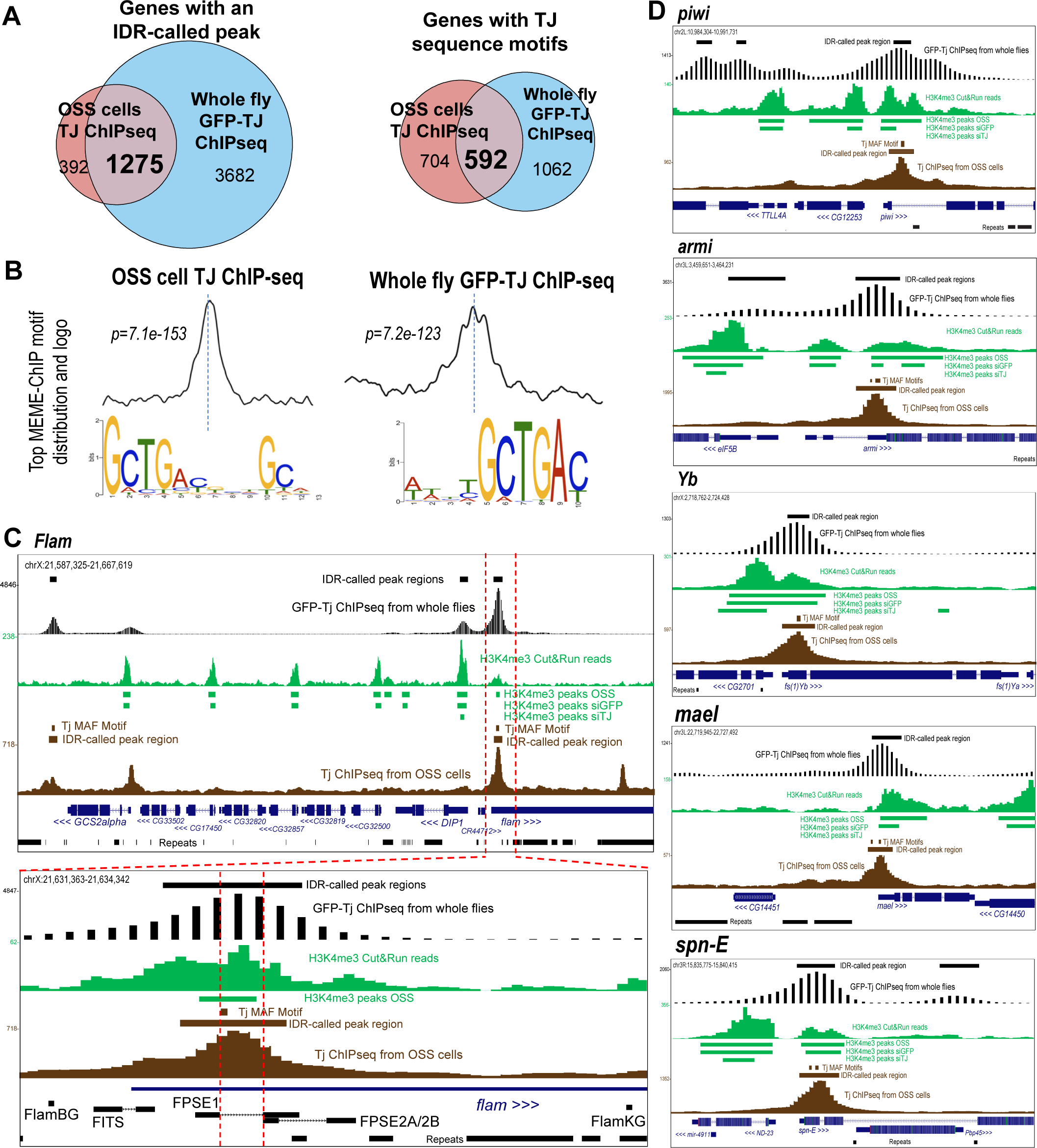
TJ binds the enhancers of *Flam* and multiple Piwi pathway genes to drive piRNA production. (A) Venn diagrams of the number of genes with IDR-called peaks from the whole flies GFP-TJ ChIPseq and the OSS cell TJ ChIPseq (left) and the genes with TJ motifs (right). (B) MEME-ChIP analysis of the top enriched motif sequence from the input of the IDR-called peaks from the OSS cell TJ ChIP-seq and the Whole fly GFP-TJ ChIP-seq datasets. Below each motif signature distribution at the center line is the sequence logo that is consistent with in vitro determined motifs of MAF transcription factors. (C) Genome browser tracks of TJ ChIP-seq and Cut&Run profiling of H3K4me3 chromatin marks at the *Flam* piRNA cluster locus. The lower panel zooms in on the *Flam* TSS. (D) Genome browser tracks at five Piwi pathway protein-coding genes with downregulated RNA expression after siTj knockdown in OSS cells.

We expected the TJ ChIPseq data from OSS cells to more-specifically reflect patterns in the ovarian follicle cells, but the GFP-TJ ChIPseq from whole flies still provided excellent independent validation of shared binding patterns as a proxy of TJ in the fly. For instance, when both OSS TJ and Whole fly GFP-TJ peaks were analyzed with the MEME program, both datasets yielded essentially the same top sequence motif with the sequence “GCTGAC” as a core motif (Fig. 5B). Remarkably, this TJ motif we determined in *Drosophila* cells matched exactly with the same TJ motif first predicted by the Fly Factor Survey that employed a bacterial 1-hybrid approach (Zhu et al. 2011) (**Supplemental Figure S6A**).

Although only ∼33% of the genes with GFP-TJ peaks contained this TJ motif, the clear majority (>77%) of the OSS cell genes with TJ peaks displayed the TJ motif (Fig. 5A), re-enforcing our confidence in focusing on the TJ-bound peaks in OSS cells. A metagene and functional annotation analysis of the distribution of TJ binding motifs shows a notable preference of TJ binding at gene TSS’s and downstream of the TSS in mostly enhancer elements versus a lower proportion on upstream promoter elements (Supp. Fig. S6B,C). However, the highly compact juxtaposition of *Drosophila* genes also biases this analysis because we could not extend the upstream region of one gene longer than ∼400bp before hitting the next gene. The majority (>81%) of 1296 genes in OSS cells have between 1-3 TJ binding motifs matching the TJ binding peaks (Supp. Fig. S6D).

In *D. melanogaster*, eight protein coding genes are configured within a ∼27 kb proximity to the *Flam* TSS, and we could define seven reproducible H3K4me3 peaks demarcating a proxy for RNA Polymerase II residence at these genes’ TSS (Fig. 5C). Six of these H3K4me3 peaks were all lost in the siTJ knockdown in OSS cells, yet there were only two definitive IDR-called TJ peaks in this region, one peak flanking the end of the upstream protein coding genes, and the other TJ peak directly overlapping the *FPSE1* deleted region in the fly mutant (Fig. 5C, lower panel). In fact, the *FPSE1* deleted region contains the TJ sequence motif and overlaps with the main H3K4me3 peak associated with this *Flam* enhancer that lies ∼450 bp downstream of the *Flam* TSS. The GFP-TJ IDR-called peak at this *FPSE1* region also matches the TJ peak from OSS cells.

The TJ binding peaks in GFP-TJ in whole flies and TJ in OSS cells were concordant across the Piwi pathway gene promoters that decreased in expression after siTj knockdown (Fig. 5D). The TJ peak that our data defines at the *piwi* promoter has much better resolution and is in perfect agreement with the previous ChIP-qPCR result from (Saito et al. 2009). Like *Flam*, the Piwi pathway genes’ TJ motif was downstream the genes’ TSS, and the most proximal H3K4me3 peak overlapping this TJ peak was lost after siTJ knockdown. Lastly, these H3K4me3 peak changes at TJ-regulated genes were exquisitely tight, such as the examples of the unaffected H3K4me3 peaks remaining at *CG2701* and *ND-23* after siTj knockdown, despite their gene starts being less than ∼500bp from the starts of *fs(1)Yb* and *spn-E*, respectively.

### TJ has been hijacked to activate TE expression in Drosophila ovary cells

Although one of TJ’s original function in the ovary is to promote transcriptomes required for follicle cell development, our data also provide strong evidence for TJ in activating the Piwi/piRNA pathway that serves as the host genome defense against TE expression. Normal fly ovary cells are usually safeguarded by a functional Piwi/piRNA pathway, yet multiple retroviral TEs are poised to reanimate and generate infective viral-like particles at the moment the Piwi/piRNA pathway is genetically compromised (Senti et al. 2023). This Senti et al study posits the theme of an on-going arms race and co-evolution between the selfish TEs and the somatic piRNA pathway that is exemplified by the enrichment of TE sequences in the *Flam* piRNA cluster locus to enable the most effective silencing of the most threatening and active TEs in *Drosophila* follicle cells (Senti et al. 2023).

We considered this paradigm to explain why several TEs long RNAs were diminished in OSS cells after siTJ knockdown (**Figure 6A**, Fig. 4D). Since the ChIPseq IDR peak calling method cannot be applied to TE sequences, we mapped the ChIPseq and Cut&Run reads to TE consensus sequences and looked for the highest intuitive peaks and differences in H3K9me3 and H3K4me3 levels. We observed pointed TJ and GFP-TJ peaks enriched at both of the LTRs of many retroviral-like TEs, like *gypsy*, *gypsy5*, *HMS-Beagle2*, and *gtwin* (Fig. 6B). Multiple other metaveridae TEs as well as some LINE and DNA TEs exhibited these TJ peaks at LTRs and ends that would correspond to the transcriptional promoter regions for the TEs (**Supplemental Figure S7**). Bolstering this result, TJ binding motifs were detected at the TJ ChIPseq peaks at these TE ends.

**Figure 6.**
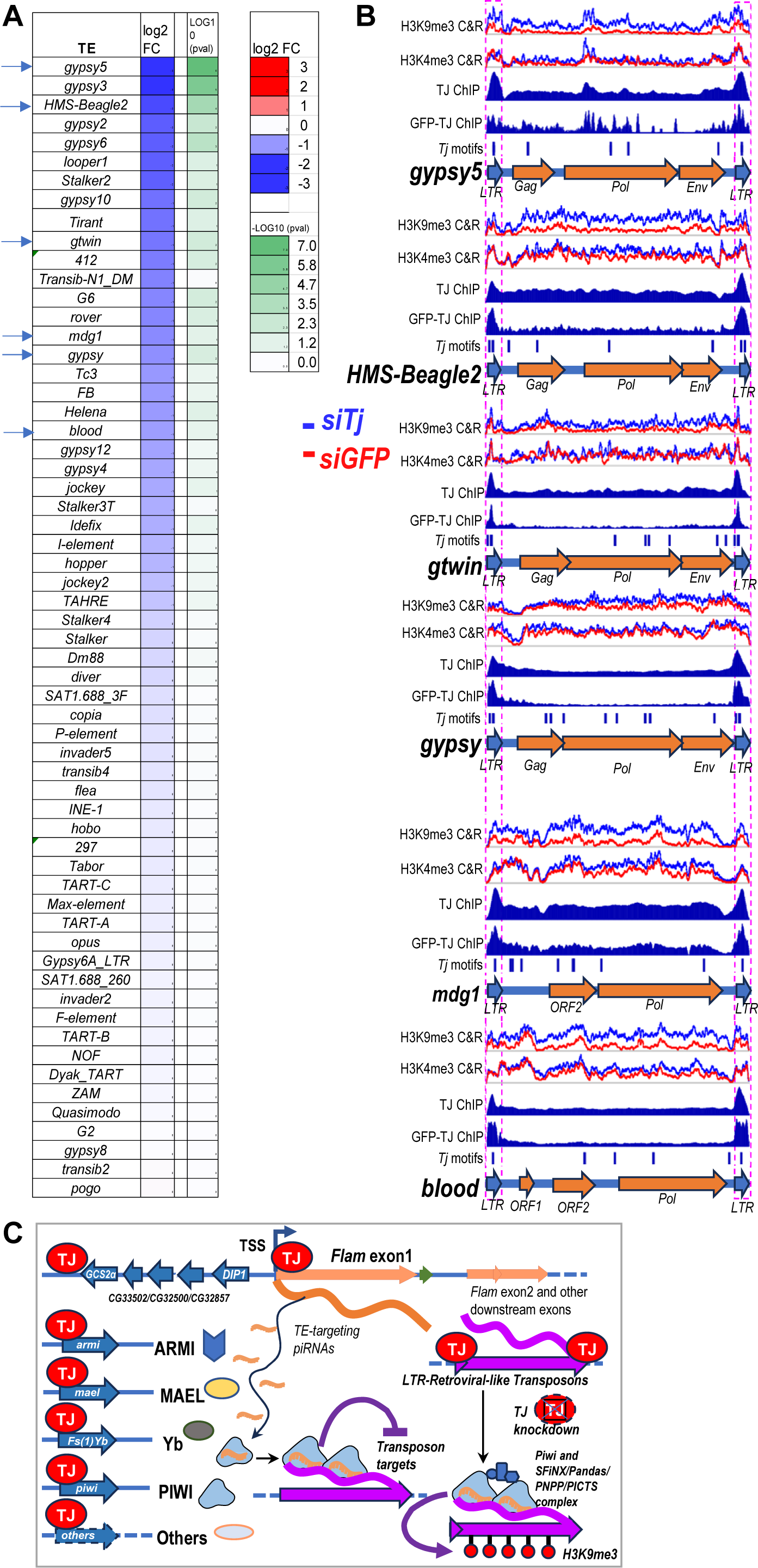
TJ activates both *Drosophila* TEs and the host TE-silencing piRNA pathway. (A) Heatmap showing lower RNA expression for certain retroviral LTR TEs after siTj knockdown. Arrows point to the TEs highlighted in the browser tracks in (B). (B) Genome browser tracks of TJ-ChIPseq and Cut&Run profiling of H3K4me3 and H3K9me3 chromatin marks at several retroviral LTR TEs actively expressed and silenced by the Piwi pathway in *Drosophila* cells and OSS cells. Additional TE browser tracks in Supp. Fig. S7. (C) Model for TJ regulating both the Piwi pathway and TEs. During siTJ knockdown, the TEs losing activation by TJ allows for residual PIWI/piRNA complexes to push more H3K9me3 silencing marks onto TE chromatin before PIWI/piRNAs also decline.

After siTJ knockdown, we observed clear increases in H3K9me3 levels at multiple TEs across much of the TE consensus’ sequences (Fig.6, Supp. Fig. S7). Since these TEs are targeted by piRNAs, this increase in H3K9me3 after siTJ knockdown may be directed by the longer residency of Piwi/piRNA complexes at TE nascent transcripts when there is not as much TJ to promote RNA Pol II transcription that would be counteracting the H3K9me3 levels. Similar to how PAF1 knockdown enhances Piwi/piRNA silencing of TE transcripts (Clark et al. 2017), TJ knockdown may operate through a similar mechanism even though the TEs’ H3K4me3 profiles did not exhibit prototypical patterns seen at the other TJ-regulated genes. Nevertheless, our data suggests TJ is also activating the expression of multiple TEs in *Drosophila* cells.

## DISCUSSION

The *Flamenco (Flam)* locus is the most prominent piRNA cluster locus expressed in *Drosophila* ovarian follicle cells that is required for female fertility and silencing of *gypsy/mdg4* TEs. Although two known *P-element*-induced mutations in the *Flam* promoter region cause loss of *Flam* piRNAs and female infertility, the *Flam* promoter and enhancer architecture was poorly understood. Our study combined promoter-bashing reporter assays, new *Flam* mutants deleting the enhancer elements, proteomics of the proteins binding these elements, and multiple transcriptomics and genomics approaches to validate the TJ transcription factor binds *FPSE1* to directly drive *Flam* expression.

Our study also uncovers a complex interplay whereby TJ activates the *Drosophila* host Piwi pathway at both protein coding and piRNA-producing genes like *Flam* to combat the LTR TEs which also leverage TJ activation (Fig. 6C). Given TJ’s high abundance in *Drosophila* follicle cells and the selfish motivations of TEs to replicate and resist host silencing mechanism, it may seem straightforward for TE promoter sequences in LTRs to evolve the TJ binding motif. To achieve détente in the TE-host genome arms race, the *Drosophila* Piwi pathway then responded by also co-evolving multiple TJ binding motifs at the protein-coding and piRNA-generating loci.

Perhaps TJ became an unwitting pawn in the TE–Piwi pathway chess game in the *Drosophila* follicle cells. Now that we have defined for the first time the thousands of genes directly bound by this large MAF transcription factor (Table S1 and S2), the main gene ontology terms linked to the TJ ChIP-seq are gonadal genesis, morphogenesis and other development processes (Supp. Fig. S6E-G). TJ is also expressed in the fly brain in a subset of neurons (Jukam et al. 2013; Konstantinides et al. 2018). Although *Tj*-null adult mutants are completely sterile from gonad collapse, the larval gonads and neurons seem to tolerate the lack of TJ, including still expressing *piwi* in *Tj*-null larval gonads (Saito et al. 2009). This suggests additional factors besides TJ in follicle cells regulates the Piwi pathway, like *ovo* for example (Alizada et al. 2024).

Will TJ’s regulation of TEs and the Piwi/piRNA pathway be conserved in other Dipterans? At evolutionary timescales, the *Flam* locus is evolving rapidly within Drosophilids and is only conserved out to *D. erecta* and *D. yakuba* that are ∼10 million years diverged from *D. melanogaster* (Malone et al. 2009; Chirn et al. 2015). The configuration of the homologous *Flam* loci in *D. erecta* and *D. yakuba* are also quite distinct from *D. melanogaster* (Malone et al. 2009; Chirn et al. 2015; van Lopik et al. 2023), and in the more closely-related *D. simulans* species the massive genomic region containing *Flam* and adjacent protein-coding gene has been duplicated (Signor et al. 2023). The *Tj* gene is conserved in Dipterans like the mosquito genomes from *Anopheles* and *Aedes*, but much further study will be needed to assess whether true *Tj* orthologs are conserved in other insects and whether the TJ binding motifs are maintained and found in the other insects’ TE sequences.

## MATERIALS AND METHODS

### Flam promoter Bashing and piRNA-targeted Luciferase reporter Assays

The pNanoLucTK plasmid (accession# KM359774) was re-engineered into pNL by replacing the TK promoter with a KpnI site for cloning various PCR-amplified genomic fragments of the *Flam* promoter/enhancer region from gDNA extracted from OSS cells. All the various *Flam* PCR fragments were generated with primers listed in Supp. Table S3, in combinations detailed in Fig. 1 and Supp. Fig. S1. PCR fragments were cloned into pNL with the NEB HiFi Assembly Kit. PCR-based mutagenesis and HiFi Assembly generated additional deletions within the longer *Flam* promoter plasmids. To further engineer the artificial intron into the *Flam* promoter reporter plasmid to make the base pNLI plastic, the artificial intron was cut out from a *Renilla* luciferase plasmid that were the *Flam-*piRNA targeted luciferase reporter plasmids previously generated in (Post et al. 2014). This artificial intron was than cloned right in front of NanoLuc at the NcoI site. All plasmids were verified by Sanger sequencing.

OSS and S2 cells were grown in T75 flasks at 25° C and harvested as pellets once fully confluent. Cells were prepared for nucleofection using SF cell line solution (Lonza). Cells resuspended in nucleofection solution were aliquoted into separate tubes with 3nmol of siRNA. Once mixed cell suspension was placed in Nucleocuvette vessel (Lonza) and nucleofection occurred in the X-unit of the 4D-Nucleofector Core Unit (Lonza) using zapping code EN-100. Nucleofected cells were placed immediately in 2X F.I.G. Media and plated in triplicates on 24-well plates at a 1:60 dilution.

Two days post-nucleofection of siRNAs, the reporter plasmid mix was prepared, per well, by diluting 0.25µg of Firefly luciferase plasmid and 0.05µg of gene specific nano luciferase reporter plasmids in 0.15M NaCl. FuGENE HD (Promega) was added to the reporter mix at 2.5µL per well and final mix was incubated for 15 minutes at room temperature. Final reporter mix was added drop wise to each well and incubated at 25° C over the next two days. Cells were lysed in 1X Passive Lysis Buffer (Promega) on a rocker for 20 minutes room temperature. Luciferase activity was measured with the ONE-Glo Stop&Glo kit (Promega) using a BioTek Synergy HT plate reader and Gen 5 program, to give the firefly luciferase expression.

### Generation of new Flam Drosophila mutants and Drosophila RNAi crosses

For the lines *FITS* and *FPSE2A/B*, Cas9 small guide RNAs and ssODNs were designed based on the *Flam* promoter bashing reporter assay results to target regions located on or upstream of the first exon of *Flam*. These oligonucleotides or their sequences were sent to Rainbow Transgenic Flies (RTF) Inc to be injected into fly embryos of the *nos-Cas9 attp2* background. When flies reached adulthood, they were crossed to *w1118* and after successful mating parents were screened for mutations by gDNA extraction. Progeny from crosses containing the mutation were collected, based on our crossing scheme, and crossed to the balancer line *Adar/FM7a*. Balancer crosses occurred over three more generations and parents were assessed for mutation by gDNA PCR for each cross. For line *FPSE1* the sequences of guides and ssODNs were consulted with RTF Inc and then sgRNA and ssODN injections, screening, and initial balancing were also conducted by RTF Inc. To keep background consistent in all lines, *FPSE1* were rebalanced over FM7a once flies were received by RTF Inc.

Several *Drosophila* RNAi crosses were prepared with different conditions as well as different Gal4 driver lines. The first set of crosses were single fly crosses consisting of a female from the line +/+; Tj-GAL4,Gypsy-LacZ/CyO; P-Tub-GAL80TS/TM6-tb, created in the Lau Lab, and a male from one of the following RNAi lines: y[1] sc[*] v[1] sev[21]; P{y[+t7.7] v[+t1.8]=TRiP.HMS01069}attP2 (RNAi-Tj#1), w; tjDf1 / CyO; UAS-tjRNAi on chr 3 (RNAi-Tj#2, doi:10.1242/dev.089896), y[1] v[1]; P{y[+t7.7] v[+t1.8]=TRiP.JF02009}attP2 (RNAi-Tj#3). Crosses were kept at 18° C until pupae began to form then they were raised to 28° C until progeny eclosed. The second set of fly crosses follow the same conditions except they were raised to 28° C during early adulthood. The final set of crosses were kept at room temperature and consist of single fly crosses between a female late-stage follicle cell GAL-4 driver, CG13083.89K-GAL4[M4]/Cyo, Dfd-eYFP (FC0), and a male from one of the previous RNAi lines. Female progeny from all crosses were collected according to the predicted knockdown phenotype and kept in vials for ∼2 days to ensure ovary development. Ovaries were then dissected and inspected under a dissecting microscope to examine knockdown efficiency by ovary morphology.

### Drosophila female fertility assays and RT-qPCR from Flam mutant ovaries

*Flam* mutant lines were assessed through an ovary squash assay and fecundity. Single sibling crosses containing either wild-type or mutant females and mutant males were set up and mated for 3-4 days at 28° C. Upon confirming successful mating flies were removed and vials placed back at 28° C. Females were carefully squashed between a microscope slide and coverslip containing green food coloring. The number of eggs per female were counted and compared between wild-type and mutants. Following this assay viable progeny from each sibling cross were counted and compared between wild-type and mutants.

For each line 20 ovaries, from both wild-type and mutant females, were dissected in cold 1X PBS. Total RNA was extracted with Monarch Total RNA Miniprep Kit (NEB) following the manufacturer’s instructions, including the DNase I treatment step. For RT-qPCR 1µg of total RNA was used for first strand cDNA preparation by using ProtoScript II Reverse Transcriptase kit (NEB) followed by LUNA Universal qPCR kit (NEB). RT-qPCR was performed in a Bio-Rad CFX Opus 96 Real-Time PCR System. Rp49 was used as the housekeeping gene to normalize downstream flamenco expression by 2-delta-delta-Ct methods.

### RNA profiling of OSS cells after siRNA Knockdown

OSS cells were grown in T75 flasks at 25° C and harvested as pellets once fully confluent. Cells were prepared for nucleofection using SF cell line solution (Lonza). Cells resuspended in nucleofection solution were aliquoted into separate tubes with 3nmol of siRNA. The mixed cell suspension was placed in Nucleocuvette vessel (Lonza) and nucleofection occurred in the X-unit of the 4D-Nucleofector Core Unit (Lonza) using zapping code EN-100. Nucleofected cells were placed immediately in 2X F.I.G. Media and plated in triplicates on 6-well plates at a 1:60 dilution. Two days after nucleofection total cell RNA was harvested using Monarch Total RNA Miniprep Kit (NEB) following the manufacturer’s instructions, including the DNase I treatment step. For RT-qPCR 1µg of total RNA was used for first strand cDNA preparation by using ProtoScript II Reverse Transcriptase kit (NEB) followed by LUNA Universal qPCR kit (NEB). RT-qPCR was performed in a Bio-Rad CFX Opus 96 Real-Time PCR System. Rp49 was used as the housekeeping gene to normalize downstream flamenco expression by 2-ΔΔCt methods.

Utilizing RNA from previous extraction from flamenco mutant ovaries and OSS knockdown cells to perform library preparations. Small RNA libraries were made using NEBNext Small RNA Library Prep Set (NEB #E7330) with up to 5µg of RNA input, and a series of terminator blocking oligos were added prior to linker ligation to suppress the 2S rRNA in the library preparation following the guidelines in (Wickersheim and Blumenstiel 2013). Library samples were amplified up to 25 total PCR cycles. Total RNA libraries were made using the Zymo-Seq RiboFree Total RNA Library Kit using up to 250ng RNA input following manufacturer’s protocol. All libraries were checked on an Agilent Bioanalyzer 2100 using either DNA 1000 or High Sensitivity DNA kit and sequenced at the BUMC Microarray and Sequencing Resource on an Illumina Next Seq 2000 using P3 flow cells for 50SE and 50PE reads for small RNA and total RNA libraries, respectively.

### Proteomics analyses of OSS cells proteins binding to FITS, FPSE1 and FPSE2 DNA segments

OSS and S2 cells were grown at 25°C in T75 flasks and harvested as 100µL pellets. Pellets were resuspended in 1X CST lysis buffer (Cell Signaling Technologies) and sonicated five times at 30% amplitude for a cycle of 15 seconds on/60 seconds off using the Q125 sonicator (QSonica). Cell lysate was spun down at 13000rpm for 10 mins at 4 °C and transferred to a new tube. Biotinylated DNA probes for FITS, FPSE1, and FPSE2 were prepared by using one normal PCR primer and one biotinylated primer designed for these specific regions. M-280 Streptavidin Dynabeads (Invitrogen) and 20pmol of biotinylated DNA probe were combined and incubated for 1 hour at 4 °C. Lysate was pre-cleared by incubating 1mg of Streptavidin Magnetic Beads (NEB) on a rotator at room temperature for 30 minutes.

After pre-clear lysate was incubated on ice with 200µg of Salmon sperm DNA (Sigma) for 5 minutes. This was followed by incubating lysate with biotinylated DNA probe for 1 hour at room temperature on a rotator. Once this final incubation was completed beads were washed three times in wash buffer followed by three washes in 1X PBS. Beads were resuspended in 100µL of 1X PBS and 80µL were sent off to the Center for Network Systems Biology (Boston University) for Mass Spectrometry while the remaining 20µL were analyzed by SDS-PAGE followed by silver staining with the Pierce Silver Stain for Mass Spectrometry kit (Thermo Scientific).

To prepare the DNA-pulled down proteins immobilized on streptavidin beads for mass spectrometry analysis, the samples were eluted by on-bead digestion in 40 ul digestion buffer (0.6M GuHCl, 4 mM Chloroacetamide, 1 mM TCEP). 0.75 ug trypsin was added to each replicate and trypsinization was carried out overnight at 37 C. Samples were acidified with 0.1% TFA and the tryptic peptides were desalted before being quantified by BCA protein assay (Pierce). Peptides from each replicate were TMT-labelled (ThermoFisher Scientific) as per the manufacturer’s protocol. Pooled TMT labelled peptides were desalted and fractionated off-line via high pH reversed-phase chromatography to obtain five fractions using different concentrations of acetonitrile (5%, 10%, 15% 20% and 50%). The peptides were dried in a SpeedVac.

The dried TMT-labelled samples were resuspended in mobile phase A (0.1% Formic acid and 2% acetonitrile) for analysis on the ThermoFisher Exploris 480 instrument interfaced to the Easy nanoLC 1200 HPLC system (ThermoFisher Scientific). The Exploris 480 mass spectrometer was operated with the FAIMS Pro interface using three compensation voltages (CV= −50, −57 and −64). The samples were loaded onto a reversed-phase C18 column (75 mm i.d. × 2 cm, Acclaim PepMap100 C18 3 mm, 100 Å, ThermoScientific) in mobile phase A and resolved over a 120-minute gradient of mobile phase B (0.1% Formic acid, 80% acetonitrile) using an EASY-Spray column (ES803A, Thermo Scientific) maintained at 45° C. The instrument was operated in Data-Dependent mode (DDA) using an MS1 resolution of 60,000 and a scan range of 350-1400 m/z with a normalized AGC target of 300% (3e6 ions) and an injection time set to auto mode. MS2 scans were acquired at a resolution of 45,000 via HCD fragmentation with a normalized AGC target of 100% (1e5 ions) and an injection time of 80 ms. The duty cycle time was set to 0.8 seconds.

Bioinformatics analysis of the raw MS files started with splitting the channels into the respective individual CV files and analyzed by MaxQuant (Ver. 1.6.7.0) that integrates the Andromeda search engine. The raw data were searched against the *Drosophila melanogaster* database from UniProt (downloaded on 11/8/22). Enzyme specificity was set to Trypsin and up to 2 missed cleavages were allowed. Cysteine carbamidomethylation was specified as fixed modification whereas oxidation and N-terminal acetylation were set as variable modifications. Protein and peptide identifications were filtered at 1% FDR using the target-decoy strategy (Elias and Gygi 2007). Peptide precursor ions were searched with a maximum mass deviation of 6 ppm and fragment ions with a maximum mass deviation of 20 ppm. The MaxQuant output file designated “ProteinGroups.txt” was used for normalization and further bioinformatic analyses. Subsequent bioinformatic analyses was performed in R. The protein group intensities identified were Log-transformed and normalized by Loess. Differential analysis was performed using the Limma R package (Ritchie et al. 2015). Moderated t tests were corrected with the Benjamini-Hochberg method for false discovery rate (FDR).

### Western Blots of Traffic Jam and other Piwi-pathway proteins

OSS cells were nucleofected to include siRNAs and then plated on 6-well plates to grow at 25°C. Once confluent, cells were harvested as pellets and resuspended in 2X SDS Sample Buffer. They were then incubated at 100°C for 5 minutes. Denatured samples were then run on a 10% mini-PROTEAN TGX Gel (Biorad) at 120V for 1 hour and 5 minutes. Using the Trans-Blot Turbo Transfer System and RTA Transfer Kits (Biorad) the proteins were then transferred onto a PVDF membrane following the quick start guide provided and the mini-TGX protocol on the system. After transfer membranes were blocked with 5% non-fat milk for 1 hour at room temperature. Next primary antibodies were prepared at a 1:5000 dilution in 5% non-fat milk and incubated with blots overnight at 4°C.

Primary antibodies specific to Traffic Jam (TJ) were generated in rabbits immunized against TJ peptide, KMEDPTINDTYVQEFD-Cys by Pacific Immunology Corp. The antibodies were then affinity purified by using SulfoLink Coupling Resin Columns (Thermo Scientific). The following day blots were washed for 5 minutes three times with cold 1X PBS. Next secondary antibodies were diluted 1:5000 and incubated with blots for 2 hours at room temperature. After incubation blots were washed for 5 minutes 3 times with cold 1X PBS. Blots were then imaged using the Odyssey CLx Imager (Li-Cor).

### Cut&Run of H3K9me3 and H3K4m3 marks in OSS cells

OSS cells were grown in T75 flasks at 25°C and some OSS cells also were nucleofected with siRNAs to create siRNA Knockdown samples. Once cells were fully confluent, they were fixed with 1mM 3,3-Dithiodipropionic acid di(N-hydroxysuccinimide ester) or DSP (Sigma-Aldrich) for 30 minutes at room temperature. After fixation DSP was quenched with 110mM Glycine and cells were harvested. Samples then were prepared using the CUTANA ChIC/ CUT&Run Kit Version 3 (Epicypher) with the following changes and additions: additional antibodies used up to 0.5µg were H3K9me3 Polyclonal (Invitrogen), TJ39/40, & TJ5 (Godt). During digest with pAG-Mnase samples were incubated for 1 hour at 4°C. after targeted chromatin digestion and release an additional reverse cross-linking step which calls for 0.1% SDS and 0.2mg/mL Proteinase K to be added to samples and then incubated at 70°C for 10 minutes. After samples cross-links were reversed up to 5 ng was used to prepare libraries with the CUT&RUN Library Prep Kit Version 1.1 (Epicypher). Library preps were sequenced at the BUMC Microarray and Sequencing Resource on an Illumina Next Seq 2000 using 50×50bp paired end sequencing on a P2 flow cell for 100 cycles.

### TJ ChIP-Seq from OSS cells

OSS cells were grown in T75 flasks at 25°C and fixed with 2% p-formaldehyde in 1X PBS for 10 minutes at room temperature. Cells were then quenched with final concentration 0.125M glycine and harvested using a cell scrapper. Cells were pelleted and washed two times with cold 1X PBS +1% BSA. Next the cell pellet was resuspended in 500µL of MNase Lysis Buffer supplemented with 1mM DTT and 0.2% SDS. Cell lysate was then transferred to Micro Tube TPX Plastic tubes (diagenode) and sonicated with medium wavelength for 10 cycles of 15 seconds on/ 30 seconds off using the Bioruptor UCD-200 (diagenode). Samples were moved back to normal tubes and cell debris was pelleted. Supernatant was moved into a new tube and brought up to 2mL with ChIP-Dilution Buffer.

Supernatant was divided evenly and incubated at 4° C overnight with 2µg of antibody. The following day 40µL of Protein A magnetic beads (Dynabead) were prepped by washing once with 1X PBS and twice with ChIP Dilution Buffer. After washes beads were mixed with supernatant and incubated for 4 hours 4°C. Once incubation was complete the following washes were performed: twice in ChIP Dilution Buffer, once in 1X TE, five times with ChIP-RIPA Buffer, twice with ChIP-RIPA/500 Buffer, twice with ChIP-LiCl Buffer, and last two washes with 1X TE. After all washes samples were eluted with 50µL 1X Elution Buffer and incubated overnight at 65°C to reverse cross-links.

On the third day elution was removed from beads and 20µg of RNase A was added to each sample then incubated at 37° C for 30 minutes. Next 100µg of Proteinase K was added to samples and incubated for 90 minutes at 37° C. After incubations samples were purified using the Monarch DNA/PCR Cleanup Kit (NEB), performing an extra wash during washing steps. Samples were eluted out in 50µL of nuclease-free water. Proceeded to DNA library prep using 25ng of input and following the standard protocol for the NEBNext Ultra II DNA Library Prep Kit for Illumina (NEB). During the PCR amplification, for target enrichment, all samples were run for only 9 cycles. All libraries were checked on an Agilent Bioanalyzer 2100 using either DNA 1000 or High Sensitivity DNA kit and sequenced at Novogene using Illumina Novaseq on a XPlus 10B flowcell.

### Bioinformatics analysis of long RNA and small RNA differential expression

Small RNA libraries were sequenced as 50nt SE reads. Some libraries were sequenced multiple times to ensure sufficient sequencing depth of >5 million reads. Files from the same library were merged. Sequencing data was checked using FastQC (Andrews 2010). Small RNA samples were run through the Mosquito Small RNA Genomics (MSRG) pipeline (Ma et al. 2021; Dayama et al. 2022). Outputs from MSRG include the alignment to genes and TEs. Total RNA libraries were sequenced as 50PE reads but were processed to look like small RNA reads so that they could also be run using the MSRG pipeline. All reads were trimmed to 35nt, R1 was reverse complemented to be in the same orientation as R2, and then the reverse complemented R1 was merged with R2. The resulting FASTQ file was then run through MSRG and processed just like the small RNA libraries.

Differential expression for OSS knockdowns were done in R using DESeq2 (Love et al. 2014) on gene counts from the MSRG pipeline. The “cooksCutoff” and “independentFiltering” arguments in the results() function were set to FALSE. The piRNA cluster counting was performed from the SAM file from MSRG was filtered based on region of interest. The following coordinates were used to extract reads mapping to piRNA clusters: flamTSS|chrX:21631207-21672456, DocStalk4|chrX:21672392-21686821, flamMDG1|chrX:21687094-21704920, flamMidEnd|chrX:21705960-22193174, flamTailEnd|chrX:22193175-22430696, 20A|chrX:21521000-21557000, piwi|chr2L:10982205-10987420, tj|chr2L:19464186-19467898, brat|chr2L:19164849-19180120. Reads per million (RPM) was determined in R by counting these reads and normalizing to library size from MSRG length distribution output.

### Bioinformatics analysis of histone Cut&Run datasets and TJ ChIPseq datasets

CUT&RUN data underwent the same quality control and alignment procedures as ChIP-seq data, using FastQC (Andrews 2010) and Bowtie2 (Langmead and Salzberg 2012). However, duplicate reads were retained due to the nature of CUT&RUN, which is reasonable to generate duplicate reads. Peaks were called against IgG controls, keeping all duplicates with MACS2. H3K4me3 peaks were called using narrow peak mode, while H3K9me3 peaks were called using broad peak mode. Unlike ChIP-seq, blacklist regions were not filtered because CUT&RUN data has different characteristics, and no specific blacklist file is available. Additionally, CUT&RUN data was mapped to the *E. coli* genome to verify spike-in proportions. The calculated scale factors from this mapping were used to normalize the BigWig files.

Raw sequencing reads from TJ OSS ChIP-seq were assessed for quality using FastQC. Adapter sequences and low-quality bases were trimmed using Trimmomatic with parameters: ILLUMINACLIP: TruSeq3-PE-2.fa:2:30:10:8, LEADING:3, TRAILING:3, SLIDINGWINDOW:4:15, MINLEN:36. The cleaned reads were then aligned to the dm6 reference genome with Bowtie2 (Langmead and Salzberg 2012) using default settings. Sambamba (Tarasov et al. 2015) filtered out duplicate and unmapped reads with the criteria -F “[XS] == null and not unmapped and not duplicate”. Samtools (Li et al. 2009) was used to sort and index the uniquely mapped reads.

MACS2 (Zhang et al. 2008) identified peaks, using input samples as controls under default parameters. The consensus peak set for all three replicates was ensured by IDR (Li 2011), evaluating peak sets across true replicates, pooled-pseudo-replicates, and pseudo-self-replicates to derive the final peak set with the highest number of peaks. Finally, blacklist regions were excluded after IDR peak calling using ENCODE’s blacklist version 2 (Amemiya et al. 2019) with Bedtools (Quinlan and Hall 2010). Motifs were identified using MEME Suite (Bailey et al. 2015) for both TJ OSS ChIP-seq and GFP-TJ ChIP-seq data. Each peak was trimmed to 500 base pairs in total, centered on the peak summit with 250 base pairs extending to each end. The most significant motif was then searched within these 500 bp peak sequences using FIMO (Grant et al. 2011) with a p-value threshold of < 5e-4 (default is p-value < 1e-4).

GFP-TJ chip seq bam files and the optimal IDR thresholded peaks were processed from the modERN resource (Kudron et al. 2018). The Whole Fly GFP-TJ ChIP-seq dataset was downloaded from the modENCODE repository under the accessing numbers BioProject PRJNA63463, Accession GSE256732. The source fly for this experiment is located at the Bloomington Drosophila Stock Center BDSC as strain #66391, and it has the genotype: w[1118]; PBac{y[+mDint2] w[+mC]=tj-GFP.FPTB}VK00033 (Chr 3, 65B2, 3L:6442676.).

Peak annotation was conducted using ChIPseeker (Yu et al. 2015; Wang et al. 2022) with tssRegion = c(−400, 600) to annotate peaks at the gene level. The reference databases used were TxDb.Dmelanogaster.UCSC.dm6.ensGene (Team BC, Maintainer BP 2019 TxDb.Dmelanogaster.UCSC.dm6.ensGene: Annotation package for TxDb object(s) and org.Dm.eg.db (org.Dm.eg.db: Genome wide annotation for Fly). Motif annotation was based on the peak annotation; if a motif was located within a peak, it was assigned to the corresponding gene. Gene enrichment analysis was performed with DAVID (Sherman et al. 2022) and visualized with tree plots using the rrvgo package (Sayols 2023).

For siTJ/siGFP H3K4me3 and H3K9me3 CUT&RUN data, reads were aligned to TE consensus sequences using Bowtie2. The aligned reads were sorted and indexed with Samtools. Scaling factors mentioned in cut and run processing section were applied for BigWig files visualization. TJ OSS ChIP-seq and TJ GFP-tag ChIP-seq reads were also aligned to TE consensus sequences using Bowtie2. The three replicates for each condition were merged into a single BAM file. These aligned reads were sorted and indexed using Samtools. siTJ/siGFP RNA-seq reads were aligned to TE consensus sequences using STAR (Dobin et al. 2013). TE counts for each sample were calculated using Samtools idxstats. TJ motif patterns and position weight matrices were downloaded from the Fly Factor Survey (Zhu et al. 2011). FIMO was used to search the motif through all TE sequences with a p-value threshold of < 5e-4.

## Resource availability

### Lead contact

Further information and requests for resources and reagents should be directed to and will be fulfilled by the lead contact, Nelson Lau.

## Materials availability

All unique/stable reagents generated in this study are available from the lead contact. Material transfer agreements with Boston University may apply.

## Data and code availability

All sequencing data produced and generated by this study is available on Sequencing Read Archive (SRA) under BioProject PRJNA1133762. See Tables S4 for specific BioSample and SRA accessions. The MSRG pipeline code can be found on the Github repository: https://github.com/laulabbumc/MosquitoSmallRNA. The mass spectrometry proteomics data have been deposited to the ProteomeXchange Consortium (Deutsch et al. 2023) via the PRIDE partner repository with the dataset identifier PXD052946 and 10.6019/PXD052946”.

## ACKNOWLEDGEMENTS

We thank Zeba Wunderlich for comments on this manuscript and Matt Layne for the use of the plate reader. We acknowledge fly husbandry assistance from Jasmine Pierre and William Dorst. We thank Dorothea Godt for various fly mutants and an aliquot of the anti-TJ guinea pig antibody. We thank Kim McCall and Jianjun Sun for GAL4 driver flies and advice. We thank Ben Czech Nicholson and Emilie Brasset for sharing previews of their data on a complementary study to our study presented here and for pointing out the modENCODE GFP-TJ ChIPseq dataset. N.C.L. is funded by NIH/NIGMS (grants R01GM135215 and equipment supplements), and a grant from the Boston University Institute of Sexual Medicine contributing support to R.K.G and J.Z.

## Author contributions

Conceptualization, N.C.L, A.R.; Methodology/Investigation, N.C.L, A.R., S.G., J.R.L., L.Y. and R.K.G.; Formal Analysis, N.C.L., A.R., S.G., J.R.L. and R.K.G.; Data Curation/Software, S.G., and J.R.L.; Writing – Original Draft, N.C.L.; Writing – Review & Editing, N.C.L., S.G., A.R. R.K.G.; Funding Acquisition, N.C.L., J.Z..

## Declaration of interests

The authors declare no competing interests.

---

**Supplemental Figure S1.**
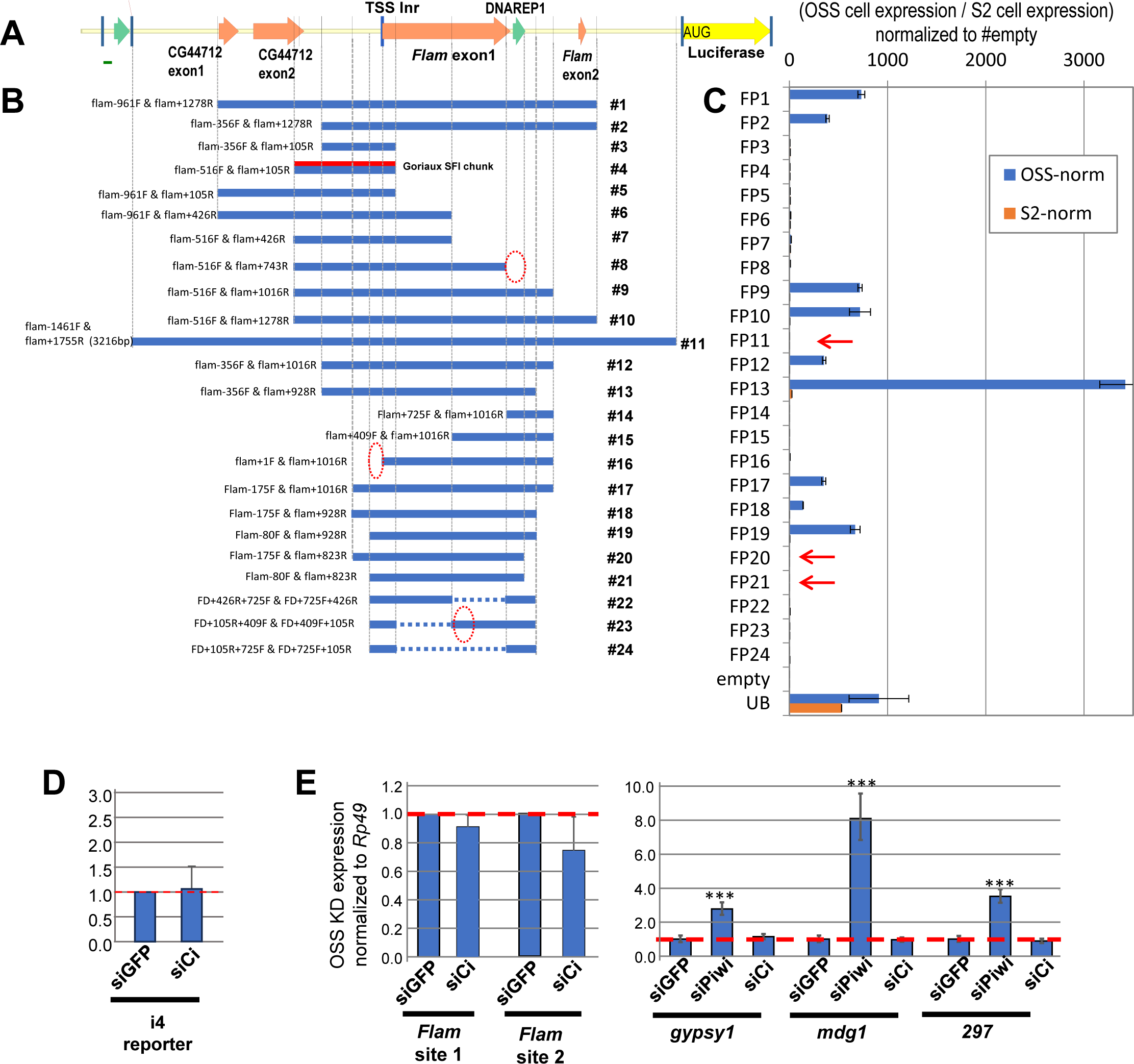
The original *Flam* promoter-bashing reporter assays lacking an intron upstream of luciferase cannot fully recapitulate the reporter activation in OSS cells. *Ci* does not impact *Flam* promoter reporters or endogenous *Flam* expression (A) Diagram of *Flam* promoter region being targeted in the promoter bashing assay format as depicted in Figure 1C except this original version lacked the additional small intron upstream of luciferase. (B) Diagram of the bashing constructs with the *FITS*, *FPSE1* and *FPSE2* regions from Figure 1 overlaid. There is an obscuring issue from the absence of a strong intron 3’ end splicing site that causes lack of luciferase expression certain constructs marked by the red arrows marked in the graph in (C) showing the corresponding normalized promoter bashing assay results from (B). (D) No change in the i4 reporter construct from Fig. 1A, the original Goriaux et al element, after siCi knockdown. (E) No change in *Flam* transcripts nor activation of TEs after siCi knockdown.

**Supplemental Figure S2.**
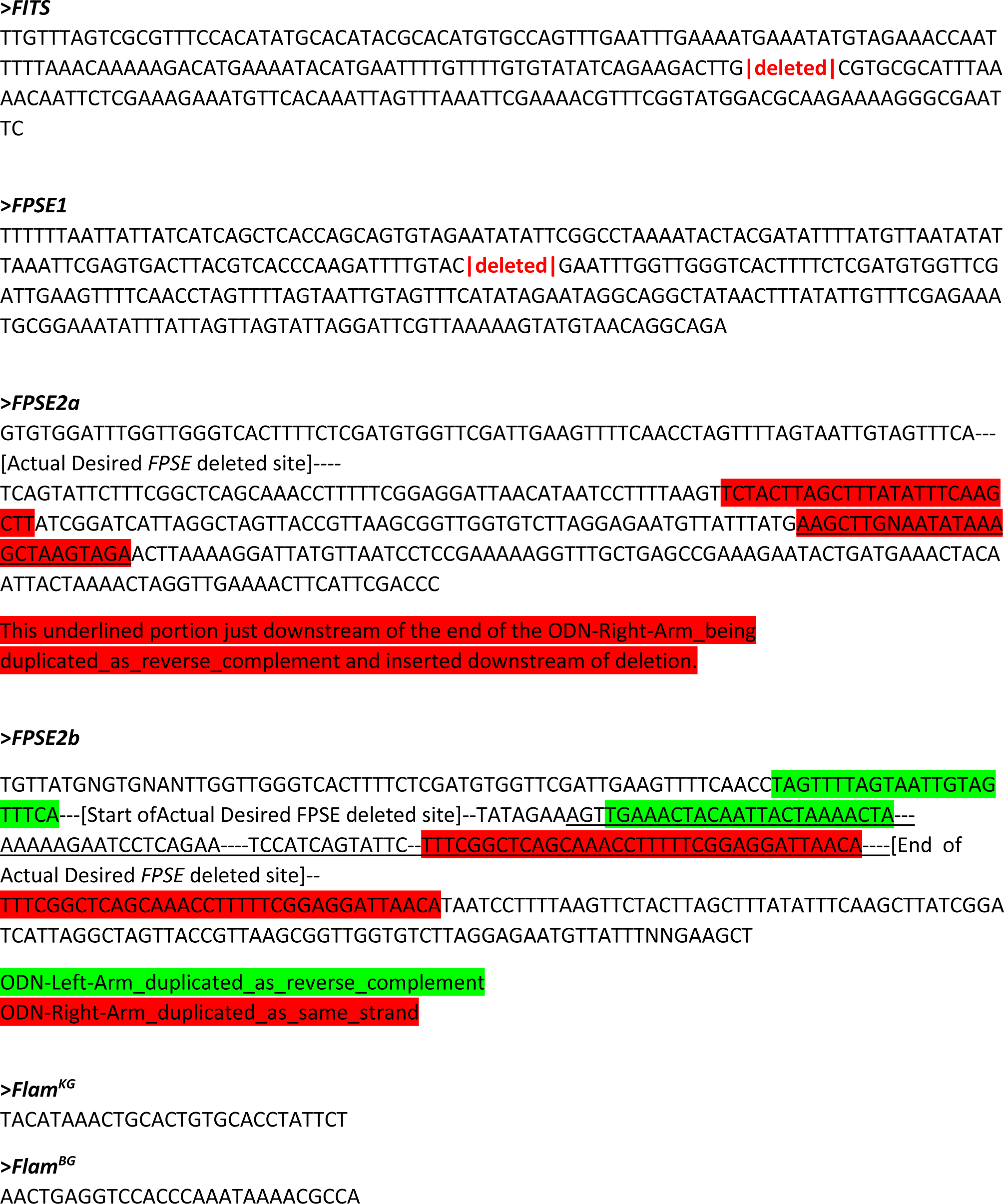
A marked-up FASTA file from the genomic PCR and Sanger sequencing of the mutated loci of the *Flam* promoter/enhancer regions in this study. The markup is mainly relevant to the determination of the portions of the ssODN repair template used to generate the *FPSE2* deletion. Unexpected ssODN sequences became integrated into the deletion site, thereby obscuring the initial mapping analysis. The short 28nt sequences for *Flam^KG^*and *Flam^BG^* demarcate flanking sequences around the *P-element* insertion in the middle of this sequence.

**Supplemental Figure S3.**
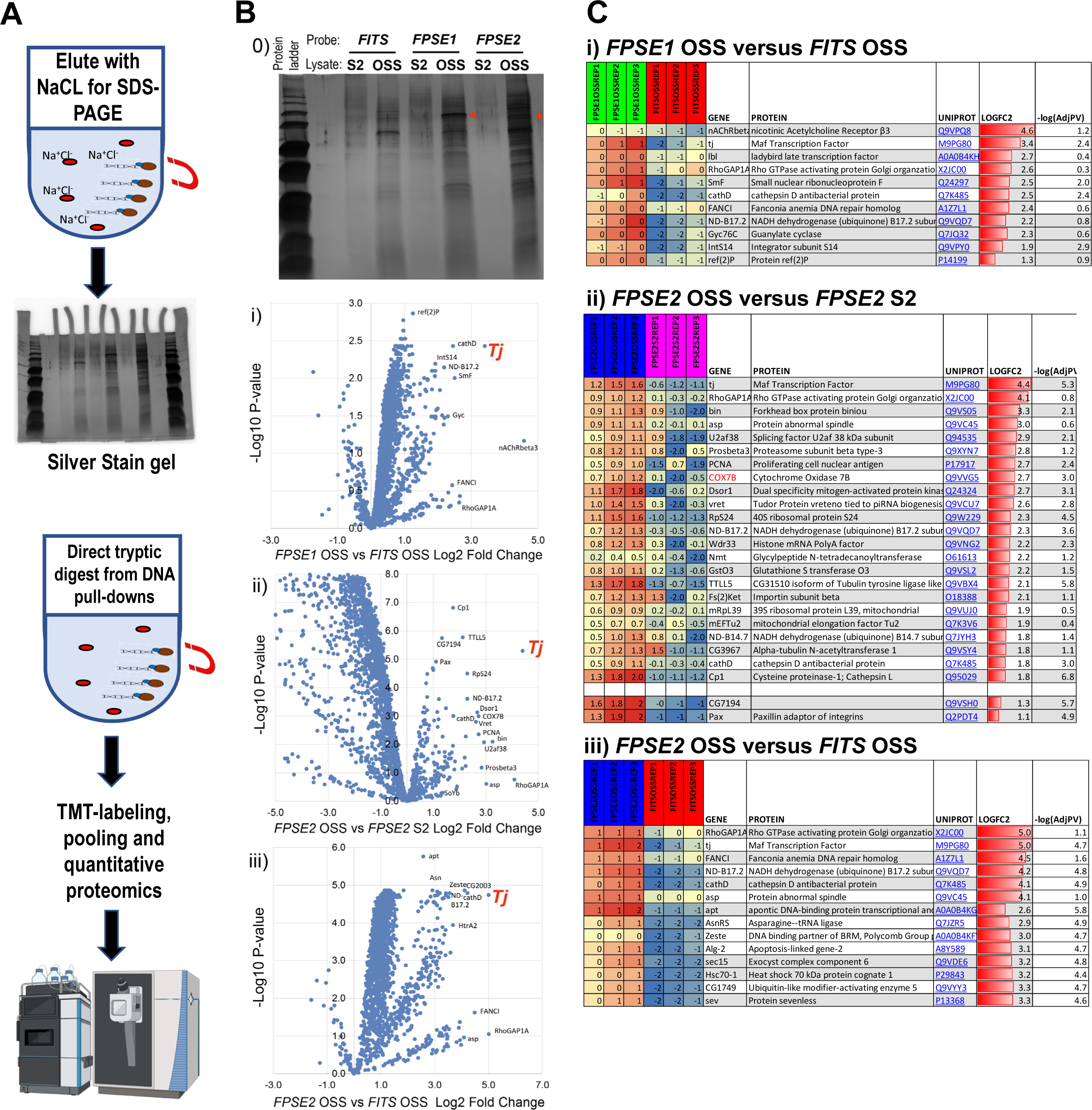
Proteomic discovery of DNA-binding protein in *Drosophil*a OSS cell lysate with biotinylated DNA probes corresponding to the *FPSE1*, *FPSE2* and *FITS* sequences. (A) Schematic representation of the quantitative proteomics approach deployed. (B) B-0 is the silver-stained SDS-PAGE visualization of novel DNA binding proteins enriched in OSS cell lysate DNA pulldowns. Red arrowheads point to the possible band of TJ. B-i, -ii, -iii are volcano plots illustrating some of the top differential proteins identified including the Traffic Jam/Tj transcription factor (red), for the indicated probes and cell lysates. (C) Table snapshots of top proteins with the greatest log2 fold difference in mass spectrometry detection differences and associated z-scores from the quantitative comparative proteomics in B-i, -ii, and -iii.

**Supplemental Figure S4.**
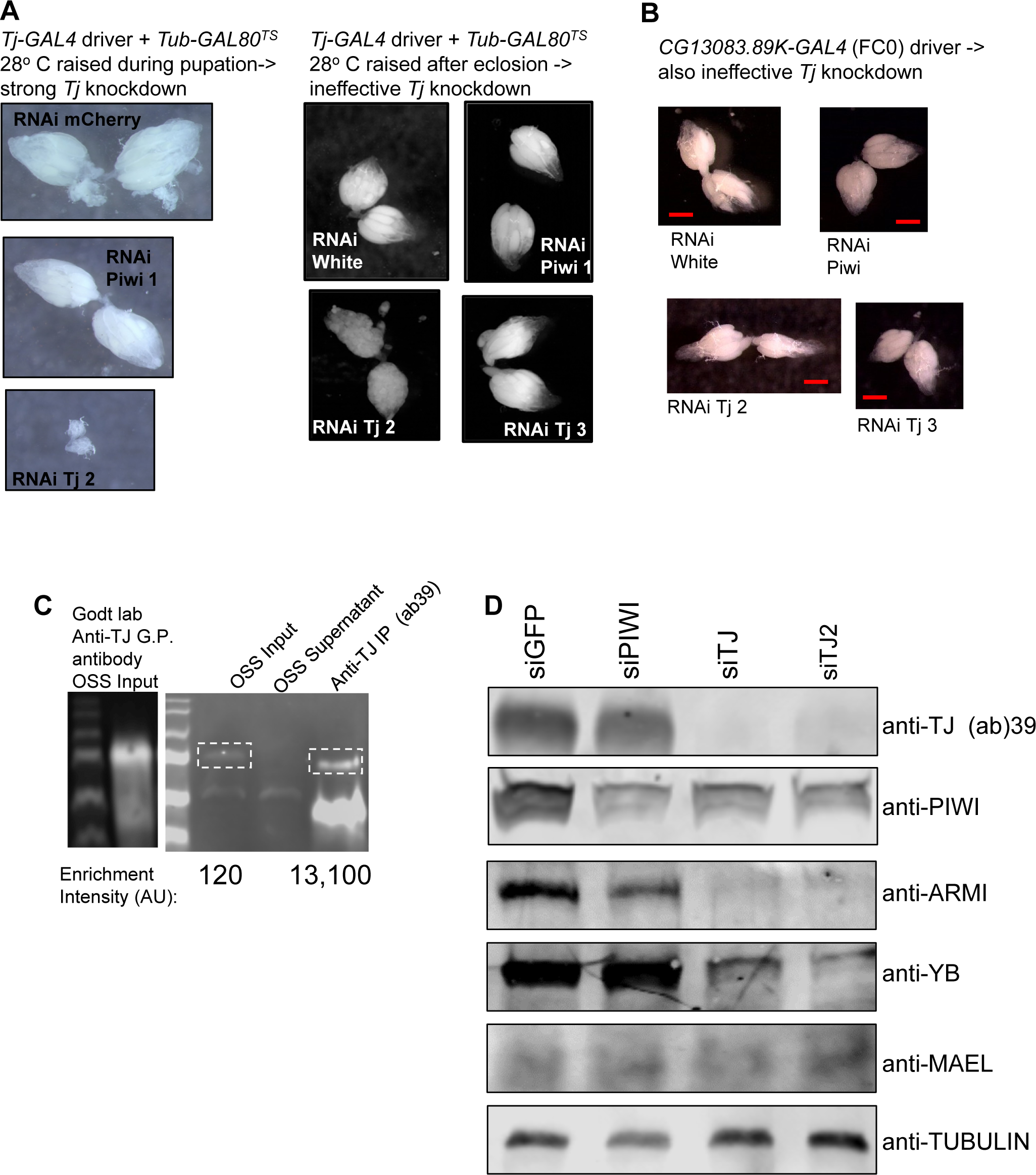
Genetic challenges with studying *Tj* knockdown in vivo and developing a new rabbit polyclonal anti-TJ antibody to perform ChIP and measure effects of siTj knockdown in OSS cells. (A) Dissected fly ovaries from progeny females from a cross of Tj-GAL4 and *Tub-GAL80^TS^* with RNAi constructs against *piwi* and *Tj*. When raising the temperature during pupation to inactivate *GAL80^TS^*, the *Tj* knockdown is effective but causes complete ovary atrophy, while raising the temperature after eclosion allows for ovary development but is ineffective at Tj knockdown. (B) A Stage 13-14 driver, *CG13083.89K-GAL4* also allowed for ovary development, but *Tj* was also ineffectively knocked down. Scale bar is 0.2mm. (C) Western blots comparing the original guinea-pig antibody to TJ from the Godt lab to our newly raised rabbit polyclonal antibody against TJ, bleed #39. OSS cell lysate used as input, and right blot shows the capacity of our antibody to immunoprecipitated TJ protein based on the enrichment intensity, AU Arbitrary Units. (D) Western blots of Piwi pathway proteins from OSS cell lysates after siRNA knockdown of *piwi* and *Tj*. Two independent siRNAs designed against the *Tj* mRNA.

**Supplemental Figure S5.**
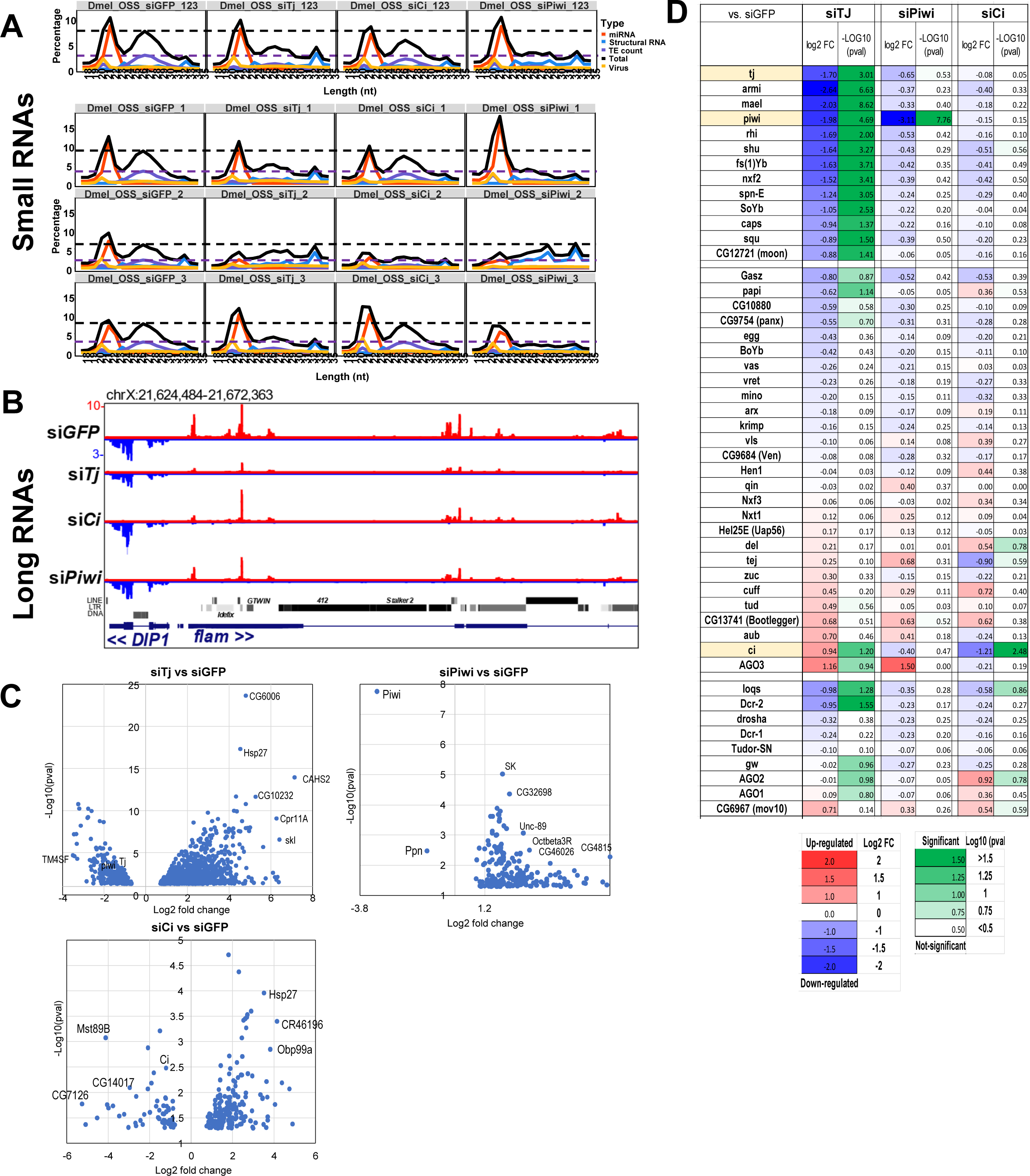
Data relating to Figure 4 whereby *Tj* knockdown in OSS cells decreases *Flam* transcripts and reduces the expression of many Piwi pathway genes. (A) Size distribution plots of small RNAs from siRNA knockdowns, showing the three individual biological replicates and the averaged combination of the three replicates (123 – top graphs). (B) Genome browser tracks showing the long RNA changes at the *Flam* locus. Note the steady expression of the DIP1 transcripts. (C) Volcano plots showing only the statistically-significant gene expression differences after a DESeq2 analysis of three biological replicates of siRNA knockdowns in OSS cells. (D) Heatmap of gene-expression changes for Piwi-pathway and

**Supplemental Figure S6.**
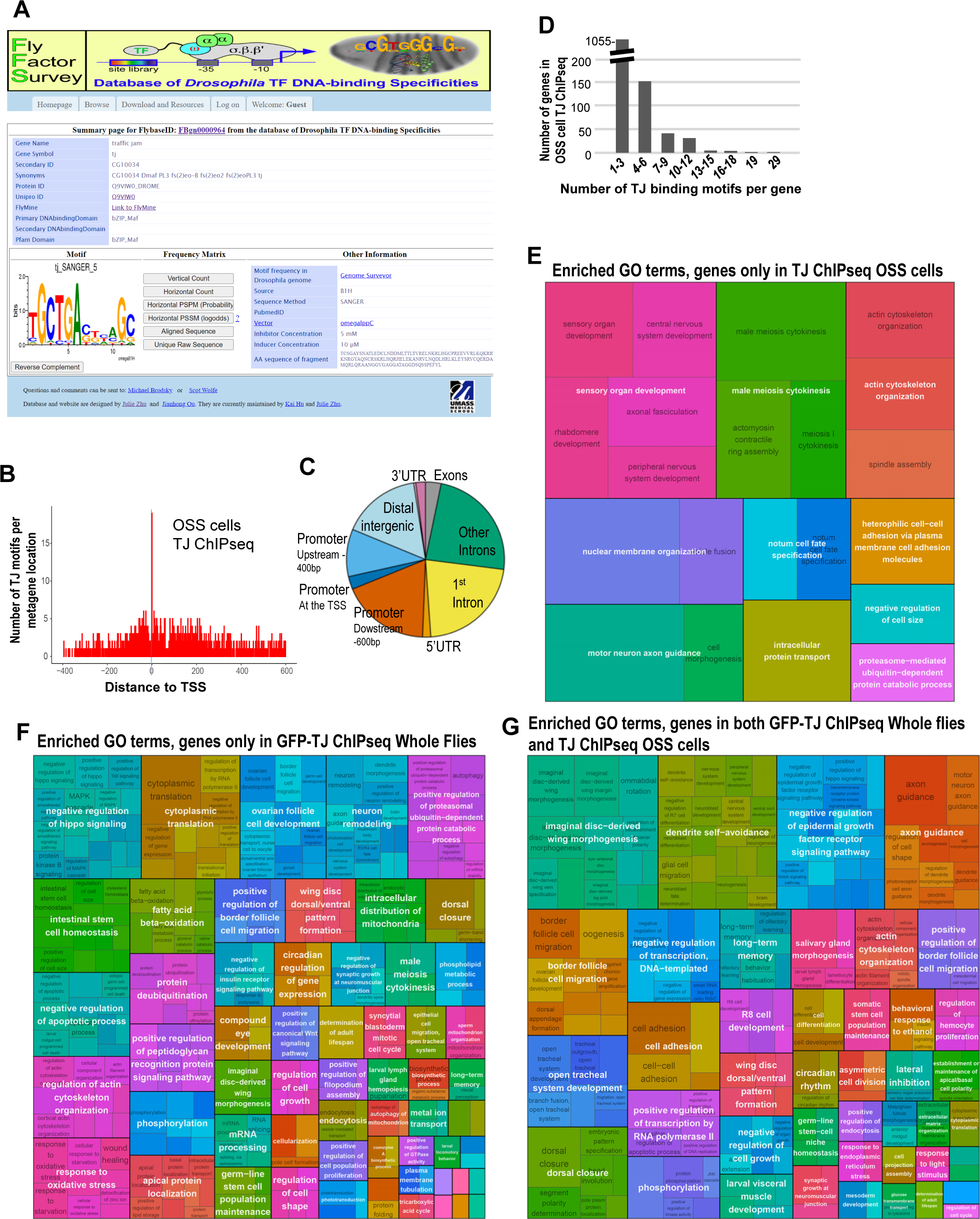
In vitro determinations of TJ binding motifs and gene ontology of *Drosophila* genes from GFP-TJ ChIPseq and TJ ChIPseq. (A) Fly Factor Survey snapshot of the Bacterial One-Hybrid Assay determination of TJ binding sequences and motif prediction. (B) Metagene plot of the distribution of the TJ motifs in the OSS cells. (C) Pie chart of the functional annotations and locations associated with the TJ motifs in OSS cells. (D) Histogram of the distribution of number of TJ motifs per gene in OSS cells. (E-G) GO term enrichment levels pictograms from genes associated just with TJ ChIP-seq in OSS cells (E), in the GFP-TJ ChIP-seq from Whole Flies (F), and for genes common to both TJ-ChIPseq datasets (G).

**Supplemental Figure S7.**
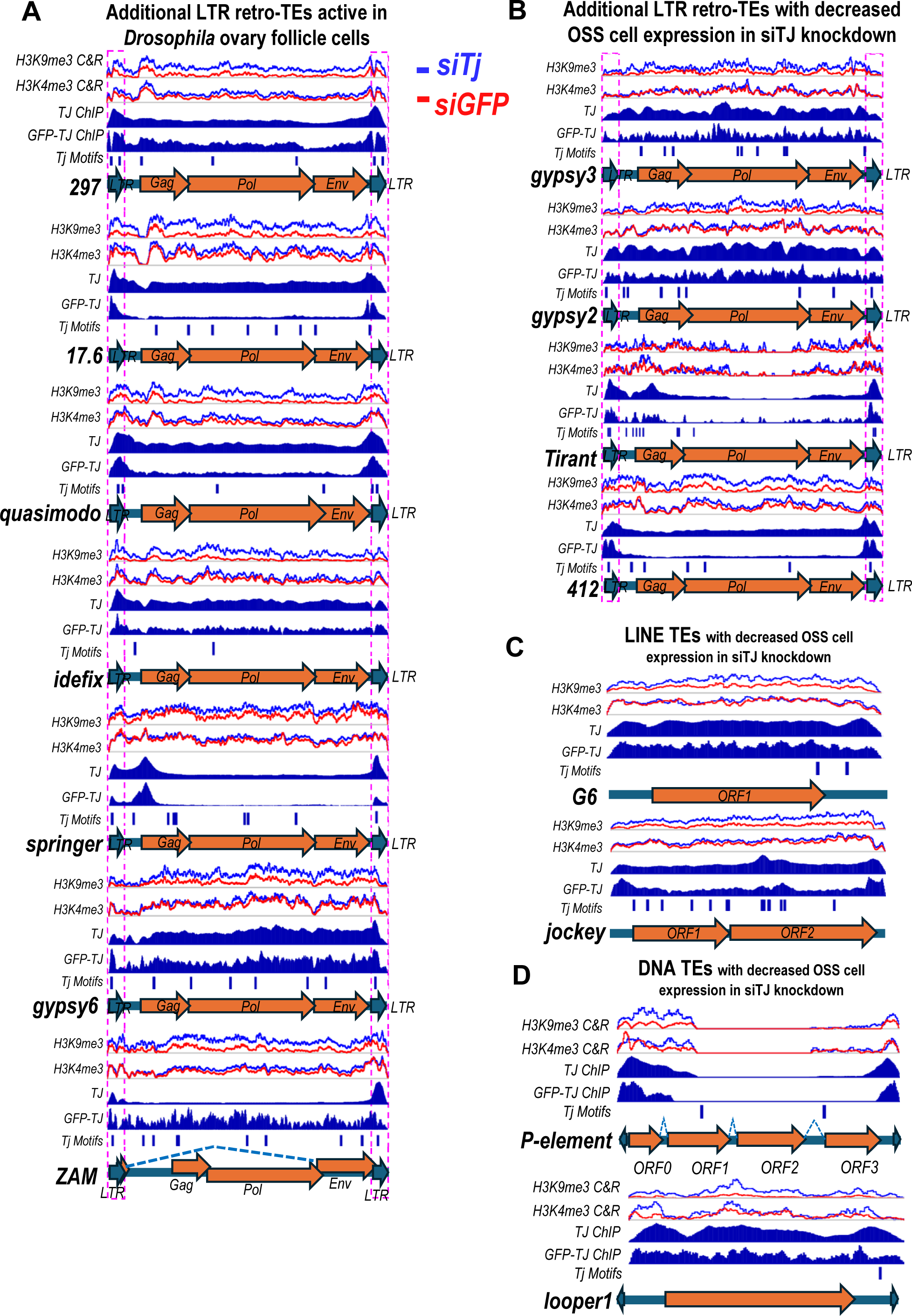
TJ ChIPseq peaks and H3K9me3 changes at other *Drosophila* TEs after siTJ knockdown. Genome browser tracks of TJ-ChIP-seq and Cut&Run profiling of H3K4me3 and H3K9me3 chromatin marks at the TEs actively expressed and silenced by the Piwi pathway in *Drosophila* cells and OSS cells in Fig. 4A. (A) Additional LTR retro-TEs active in *Drosophila* ovary follicle cells. (B) Additional LTR retro-TEs with decreased OSS cell expression in siTJ knockdown. (C) Examples of two LINE TEs with decreased OSS cell expression in siTJ knockdown. (D) Examples of two DNA TEs with decreased OSS cell expression in siTJ knockdown.

**Table S1. The list of *Drosophila* genes in OSS cells with TJ peaks and TJ binding motifs.**

**Table S2. The list of *Drosophila* genes in whole flies with GFP-TJ peaks and TJ binding motifs.**

**Table S3.**
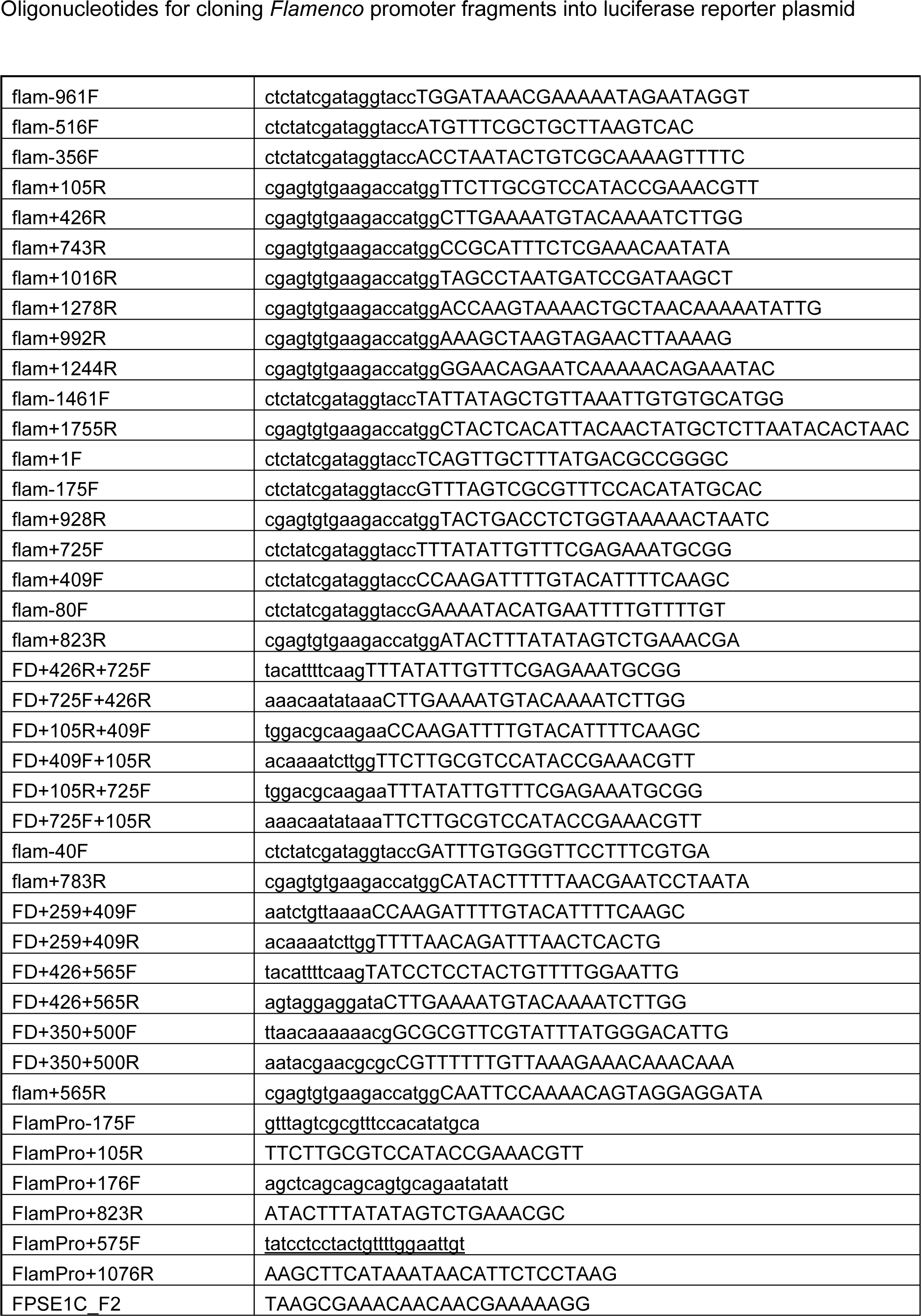
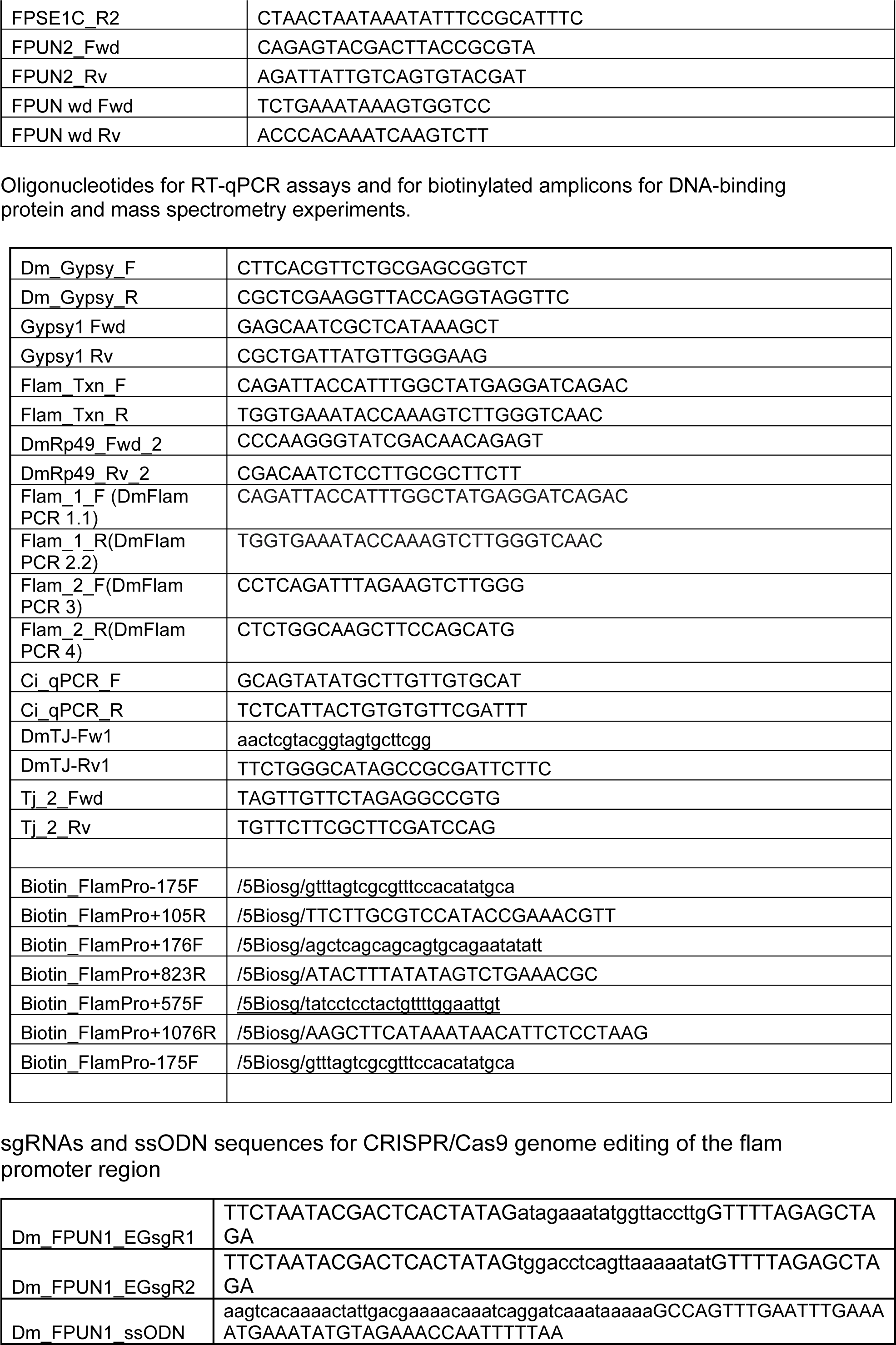
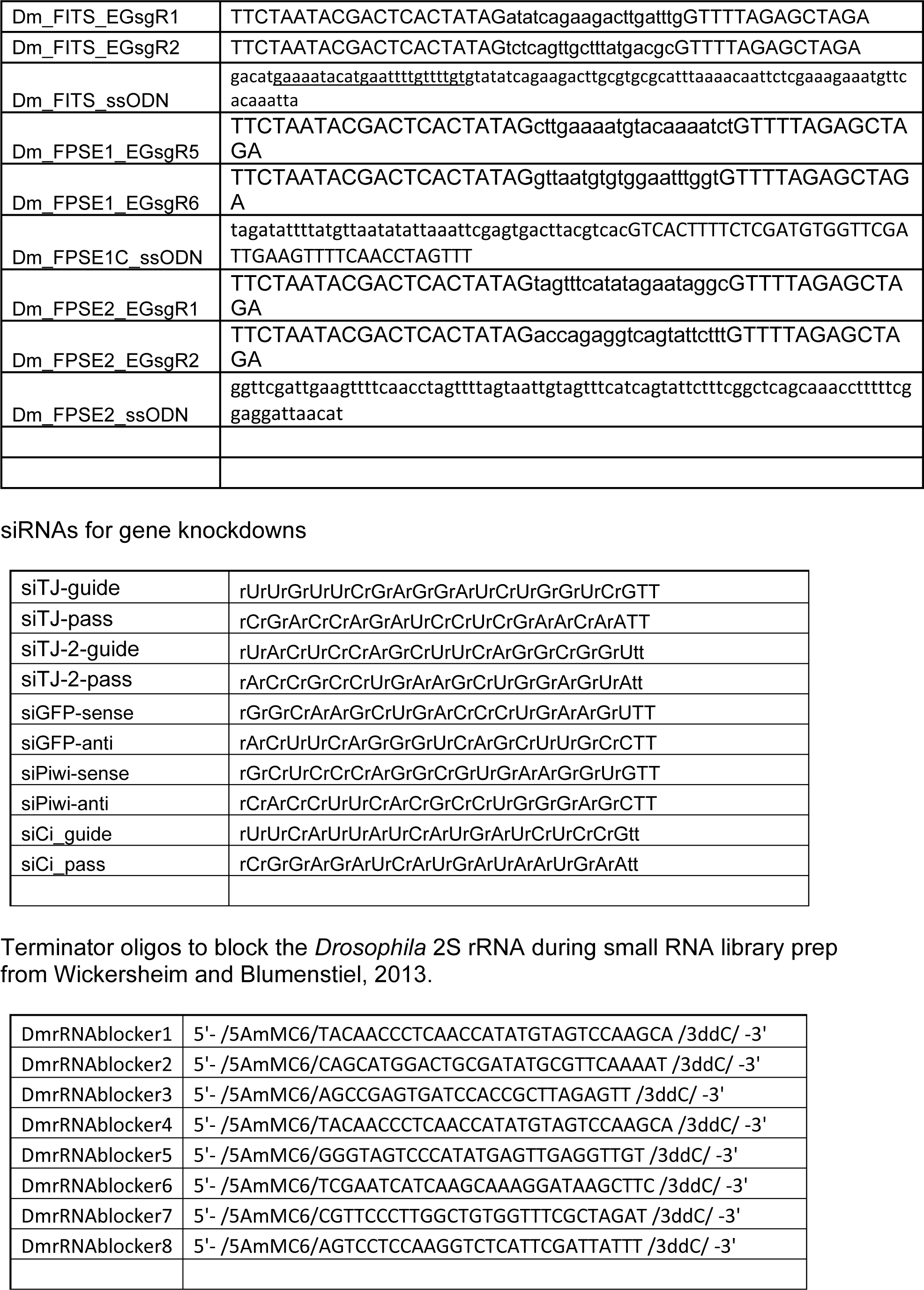
Oligonucleotide primers sequences, sgRNAs, ssODNs, and siRNAs used in this study.

**Table S4.**
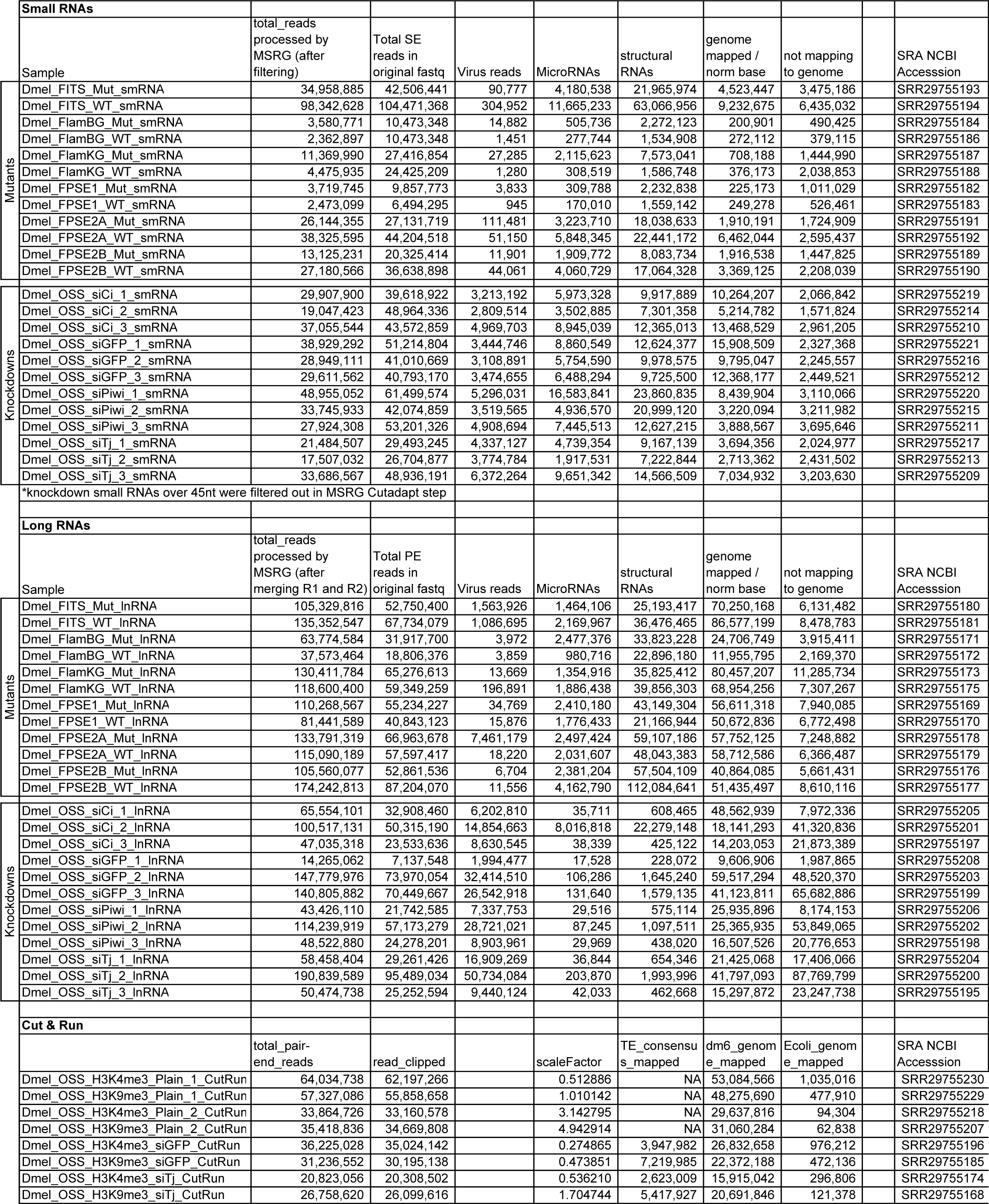
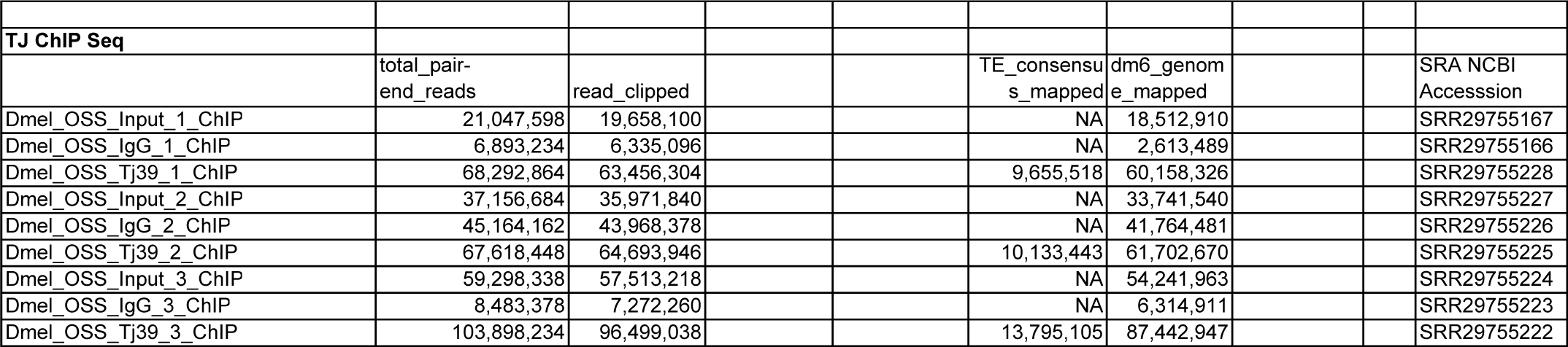
Illumina Sequencing Statistics of the smRNA, lnRNA, Cut&Run, and ChIPseq libraries generated in this study.

